# Epitranscriptomic Dysregulation in Stress-induced Psychopathologies

**DOI:** 10.1101/2021.02.17.431575

**Authors:** Dan Ohtan Wang, Kandarp Joshi, Anand Gururajan

**Author notes:** Corresponding Author. Address: Level 6, Building F, 94 Mallett Street, Camperdown NSW 2050, Australia.

## Abstract

To date, over 100 different chemical modifications to RNA have been identified. Collectively known as the epitranscriptome, these modifications function to regulate RNA stability and as such, represent another mechanistic layer of post-transcriptional gene regulation. N6-methyladenosine (m6A) is the most common RNA modification in the mammalian brain and has been implicated in a number of processes relevant to neurodevelopment, brain function and behaviour. Here, following brief descriptions on epitranscriptomic mechanisms, we will review the literature on the potential functions of the m6A-methylome in fine-tuning gene expression which include prescribing localisation of transcripts in distal compartments as well as interactions with microRNAs and long non-coding RNAs. We will then discuss findings from rodent and human studies for stress-induced disorders - major depression and post-traumatic stress disorder – which support a hypothesis for a dysregulation of the m6A-methylome and the m6A-machinery in the pathophysiology. To support this, we have included a bioinformatic analysis of publicly available single-cell RNA-sequencing and bulk transcriptomics datasets which suggests an altered m6A-methylome as a consequence of dysregulated cell- and regionally-specific expression of key enzymes involved in the ‘writing, reading and erasing’ of m6A. We hope this review will generate further interest in the field of epitranscriptomics, opening up new lines of research into its involvement in psychiatric disorders.

## Introduction

Over recent decades, more than 100 different types of RNA chemical modifications have been identified that confer RNA transcripts with information beyond the sequence of four canonical nucleotide bases, expanding the genetic vocabulary to collectively form a new layer of post-transcriptional regulation, known as the epitranscriptome. Epitranscriptomic modifications are added during or after transcription, are generally transient, site-specific, transcript-specific, and reversible which gives them the ability to modulate gene expression in a stimulus-dependent and spatiotemporal manner ^1,2^.

One of the most prevalent and highly specific epitranscriptomic modifications in the mammalian nervous system is N6-methyladenosine (m6A) ^3^. A complex assortment of nuclear and cytoplasmic enzymes is involved in the ‘writing’ and ‘erasing’ of m6A at motifs which are mainly localised to 3’UTRs near stop codons of thousands of mRNAs ^4,5^. Recent work has highlighted that the m6A-methylome in the human and mouse brain is highly specific in comparison to non-brain tissue and there are nearly 4,000 m6A-containing orthologous transcripts present in the cerebellum of both species ^3^. As with all other RNA modifications, the m6A-methylome represents a fundamental layer of regulation in the biology of the neuron such that its absence or mutation can have a significant impact on viability ^6^. Accordingly, over the last decade the m6A-methylome has been implicated in regulating gene expression programs involved in neurodevelopment, neuroplasticity, neurophysiology and behaviour ^7^.

In this review, we have firstly outlined our current understanding of the m6A enzymatic machinery – ‘writers, readers and erasers’ - and the m6A-methylome. We then examined the m6A-directed regulation of local protein translation in various intraneuronal compartments as well as interactions between the m6A-methylome/machinery and other post-transcriptional regulatory mechanisms to fine-tune gene expression. Stress can trigger immediate effects on neurotransmission, synaptic plasticity and metabolism as well as long-lasting molecular and structural changes in the brain which are underpinned by genome-wide alterations in gene expression ^8^. Since stress is also a major risk factor for psychiatric disorders such as major depression and post-traumatic stress disorder (PTSD), we then reviewed studies with animal models for aspects of these disorders as well as human studies which leads to our hypothesis that an altered m6A-methylome and the dysregulated expression of m6A-machinery is a feature of their pathophysiology.

## Writing, Reading & Erasing

### m6A Writers

The methyltransferase complex (MTC) is comprised of two core proteins - the catalytic METTL3 (methyltransferase 3) and the RNA-stabilizing METTL14 (methyltransferase 14) – together with several cofactor proteins including the scaffolding protein WTAP (Wilms tumor-1 associated protein), VIRMA (Protein virilizer homolog) and RBM15/15B (RNA binding protein) that facilitate identification of target modification sites and co-transcriptional m6A methylation as well as localisation of the MTC in nuclear speckles ^5,9,10^. Two other m6A writers include METTL16 and ZCCHC5 (retrotransposon gag-like protein 3) are not part of the canonical MTC and appear to target unique RNA subtypes ^11,12^. As mentioned above, the MTC deposits m6A at a sequence motif of DRACH (D=A/G/U, R=A/G, H=A/C/U). Notably, m6A deposition at the 3’UTR is the basis for potential interactions with microRNAs that target the 3’UTR and are themselves implicated in regulating gene expression ^3,13^. Recent studies have also shown in mammalian cells that adjacent m6A sites were prone to clustering within a 200nt region; this is evidence in support of m6A modifications interacting with each other to influence post-transcriptional processing of the same transcript ^14,15^. While m6A deposition on RNAs is mainly a nuclear process, recent work suggests that methyltransferase activities may also occur outside the nucleus where they may take on additional functions beyond ‘writing’ ^16–19^.

Factors which determine site-specific methylation at the DRACH motif include key structural features on mRNAs such as the terminal exon-exon junction near the stop codon ^20^, direct recruitment of the MTC to RNA Pol II ^21^ as well as binding sites for RBM15/RBM15B located near the m6A sites ^22^. A more critical role for METTL14 in directing site-specific methylation was identified via its interactions with the epigenetic modification, histone 3 trimethylation at Lys36 (H3K36me3). Specifically, the METTL14-H3K36me3 complex was found to direct the binding of the MTC to adjacent RNA Pol II and to deposit m6A co-transcriptionally on nascent RNAs ^23^. Interestingly, the METTL3/METTL14 complex was also found to be involved in demethylation of histone 3 lysine 9 dimethylation (H3K9me2), via interactions with the m6A ‘reader’ YTHDF1 and KDM3B (lysine-specific demethylase 3B) ^24^. These findings would suggest that complex epigenetic-epitranscriptomic interactions exert their influence on gene expression across transcriptional and post-transcriptional processes. Transcript-specific methylation may be mediated by microRNAs as well as transcription factors which interact directly with the MTC ^25^.

Knockout of *Mettl3* has been shown to be embryonic lethal ^26^ and conditional male and female knockouts of *Mettl3* had deficits in spermatogenesis and oogenesis, respectively ^27^. Conditional knockout of *Mettl3* in the developing mouse brain caused cerebellar defects ^18^. Conditional knockout in adult forebrain excitatory neurons was not found to have an impact on gross brain morphology, locomotor activity or anxiety-like behaviours but affected memory formation and increased marble burying behaviours ^28,29^. Several METTL3 inhibitor compounds have been developed but their in vivo profiles are undefined ^30^. CRISPR-Cas9 mediated knockout of *Mettl14* in mice induced embryonic lethality ^31^ and conditional knockout in the brain induced postnatal lethality ^32^. Analysis of neural precursor cells from these conditional knockouts showed a reduction in nuclear export of m6A-RNAs ^33^. Conditional deletion of *Mettl14* from dopaminergic neurons in the mouse striatum increased neuronal excitability and impaired learning and performance ^34^.

### m6A Readers

The detection and processing of m6A-RNAs is mediated by one of two processes. Direct ‘reading’ involves binding of specific RNA binding proteins (RBP) whereas indirect ‘reading’ involves alterations to the RNA structure, thereby rendering it accessible to a different set of RBPs. The YTH domain-containing family of proteins (YTHDC1/2, YTHDF1/2/3) are direct m6A ‘readers’ by YTH domain binding to m6A sites^5^. ‘Readers’ also contain low-complexity domains which facilitates their phase separation into compartments such as P-bodies or neuronal granules, particularly when bound to poly-methylated m6A-RNAs, creating a ready-to-process pool of transcripts ^35–37^.

There has been considerable debate on whether each of the m6A ‘readers’ have unique functions or whether there is redundancy in ‘reader’ function. Until recently, YTHDF1 and YTHDF2 were thought to facilitate translation and degradation, respectively, and YTHDF3 was implicated in both with each DF ortholog having affinity for subsets of m6A-RNAs ^5,38,39^. It now appears, at least in specific contexts, that all three degrade m6A-RNAs ^27,40^. Furthermore, while depletion of individual DF proteins has minimal impact on transcript abundance and stability, depletion of all three leads to a stable m6A-RNA profile, suggesting that each ortholog has the ability to compensate for the loss of another but is also dependent on its unique expression profile ^40^. So, if YTHDF1 and YTHDF3 are not involved in facilitating translation, are there any other RBP mechanisms that mediate this process? One may involve direct binding of 5’UTR m6A to the eukaryotic initiation factor 3 to initiate translation (eIF3), even in the absence of the eukaryotic initiation factor 4E which is a cap-binding protein that normally recruits eIF3 ^41^. Interestingly, a second mechanism may be with METTL3 itself acting as a translational enhancer. Once the MTC has methylated its target transcript, METTL3 does not disengage but rather stays bound to the RNA when it has been exported to the cytoplasm, where it then interacts with eIF3 to initiate translation ^42^.

In mice, YTHDC1 and YTHDC2 are reportedly necessary for fertility and embryogenesis ^43–46^. CRISPR-Cas9 mediated knockout of *Ythdf1* in mice resulted in impairments in behaviour and hippocampal synaptic plasticity ^47^. CRISPR-Cas9 mediated knockout of *Ythdf1* or *Ythdf3* in mice was not found to affect fertility or viability but knockout of *Ythdf2* severely affected both these functions; *Ythdf2* heterozygote mice required a functional copy of *Ythdf*1 or *Ythdf3* to survive ^27,48^.

A key feature of some m6A ‘readers’ is their reported ability to rapidly respond to changing stimuli by moving between compartments. For example, studies have shown that YTHDF1/2 are able to redistribute themselves to the nucleus or cytosol to mount an effective response to heat shock stress or viral infection ^49,50^. The activity of neuronal YTHDF1 in particular appears to be stimulus-dependent with low basal constitutive activity but enhanced translational activity following a depolarising stimulus or in response to injury signals ^47,51^.

Indirect readers bind to m6A-RNAs which have undergone a change in conformation, also known as an ‘m6A-structural switch.’ This change is due to the instability of the m6A·U base pair compared to A·U which results in simple, linear conformations instead of complex structures, making them more accessible to RBPs such as the nuclear fragile X mental retardation protein (FMRP) which can compete with YTHDF2 for binding and necessary for m6A-RNA transport into the cytoplasm ^33,52^.

It remains unclear if m6A ‘readers’ show any specificity towards particular m6A-sites or m6A-RNAs. They may show preference to m6A sites on certain regions of RNAs (eg. CDS vs 3’UTR), interact with other RBPs such as FMRP that share similar or overlapping consensus sites or their enrichment in structures such as nuclear speckles predisposes their activity towards specific m6A-modified transcripts ^38^. One important caveat to this discussion of direct vs indirect readers is the challenge in differentiating between the binding of an RBP directly to an m6A-RNA or binding to an m6A-RNA due to an m6A-structural switch. RNAs differ not just by methylation status but also by structure and so in interrogating the functional biology of specific readers, it would be necessary to develop sensitive methods to detect RNA structures in vivo.

Interestingly, there also appears to exist a class of so-called m6A ‘anti-readers’. These proteins are repelled from the m6A-RNA by the modification, and possibly in combination with the context RNA sequence. Two such ‘anti-readers’ are G3BP1 and LIN28A and by binding to the same site, may counteract the destabilising effects of ‘readers’ on m6A-RNAs^53, 54^.

### m6A Erasers

Fat mass and obesity associated (FTO) protein and AlkB Homolog 5, RNA Demethylase (ALKBH5) are the two main demethylases or m6A ‘erasers.’ FTO is enriched in the brain and known to localise both in the nucleus and the cytoplasm; its shuttling between these two compartments is reportedly facilitated by exportin 2 ^55^. The specificity of FTO remains equivocal as evidence suggests that it preferentially demethylates another RNA modification, N6,2’-O-dimethyladenosine (m6Am), over m6A and its absence does not significantly impact m6A stoichiometries ^28,56,57^, findings which are odds with more recent work ^58,59^. A number of global *Fto* knockout mice have been generated using different gene editing strategies resulting in a variety of phenotypes. One strategy led to *Fto* knockout mice displaying reduced anxiety and depressive-like behaviours ^60^, whereas another produced mice that had a hyperactive hypothalamic pituitary adrenal (HPA) axis, increased anxiety-like behaviours and impaired working memory ^61,62^. The latter knockouts also suffered from deficiencies in the processing of brain derived neurotrophic factor (BDNF). Conditional deletion of *Fto* in the brain impaired growth and increased metabolism ^63^; conditional deletion in adult excitatory neurons reduced marble burying behaviours ^28^. Meclofenamic acid is an FTO inhibitor and a potential tool compound to further probe its functional role ^64^. A more specific inhibitor, FB23-2, has been developed but its neuropharmacology remains unexplored ^65^.

ALKBH5 is an m6A ‘eraser,’ colocalising with nuclear speckles, enriched in reproductive tissue ^66^ but also found in the brain ^67^. In vitro work has shown that a deficiency in ALKBH5 results in increased mRNA localisation in the cytoplasm in contrast to nuclear accumulation in cells with ALKBH5 suggesting that, as a consequence of its demethylating properties, ALKBH5 also regulates mRNA export into the cytoplasm ^66^. Interestingly, recent work by Huang et al., ^68^ has shown that localisation of ALKBH5 in astrocytes in vivo is under the regulation of circular RNA, circSTAG1 (see below). Not unlike FTO, ALKBH5 also shows promiscuity in its demethylating effects on other modifications including m6_2_A in ribosomal RNA ^69^. Knockout of *Alkbh5* in male mice resulted in smaller testes and defective spermatogenesis ^66,70^.

The localisation and molecular functions of the above described ‘writers,’ ‘readers’ and ‘erasers’ have been represented in Figure 1. In table 1 we have described the reported phenotypes of relevant transgenic mice.

**Table 1.**
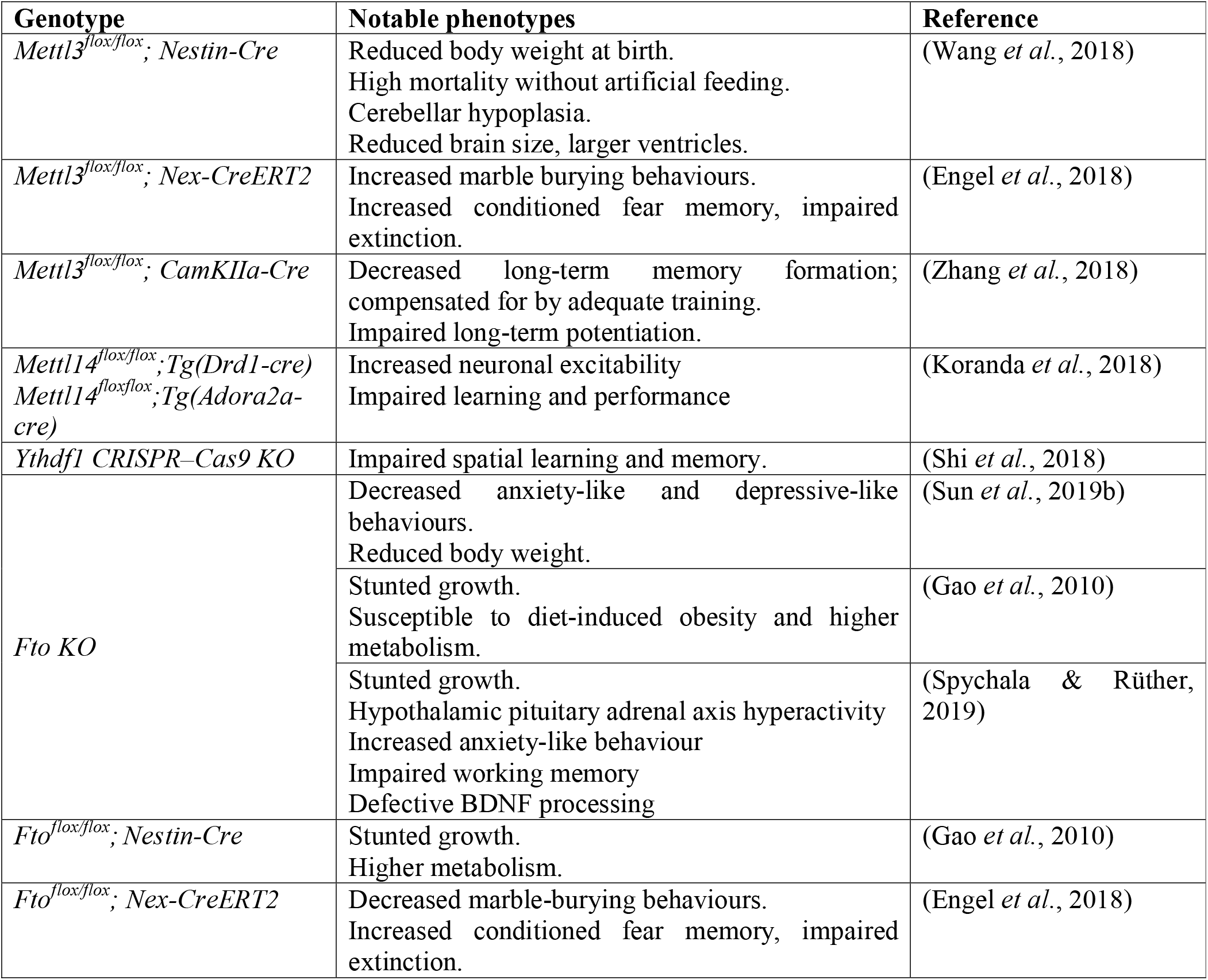
Genetically engineered mice to study the roles of m6A ‘Writers,’ ‘Readers’ and ‘Erasers.

**Figure 1.**
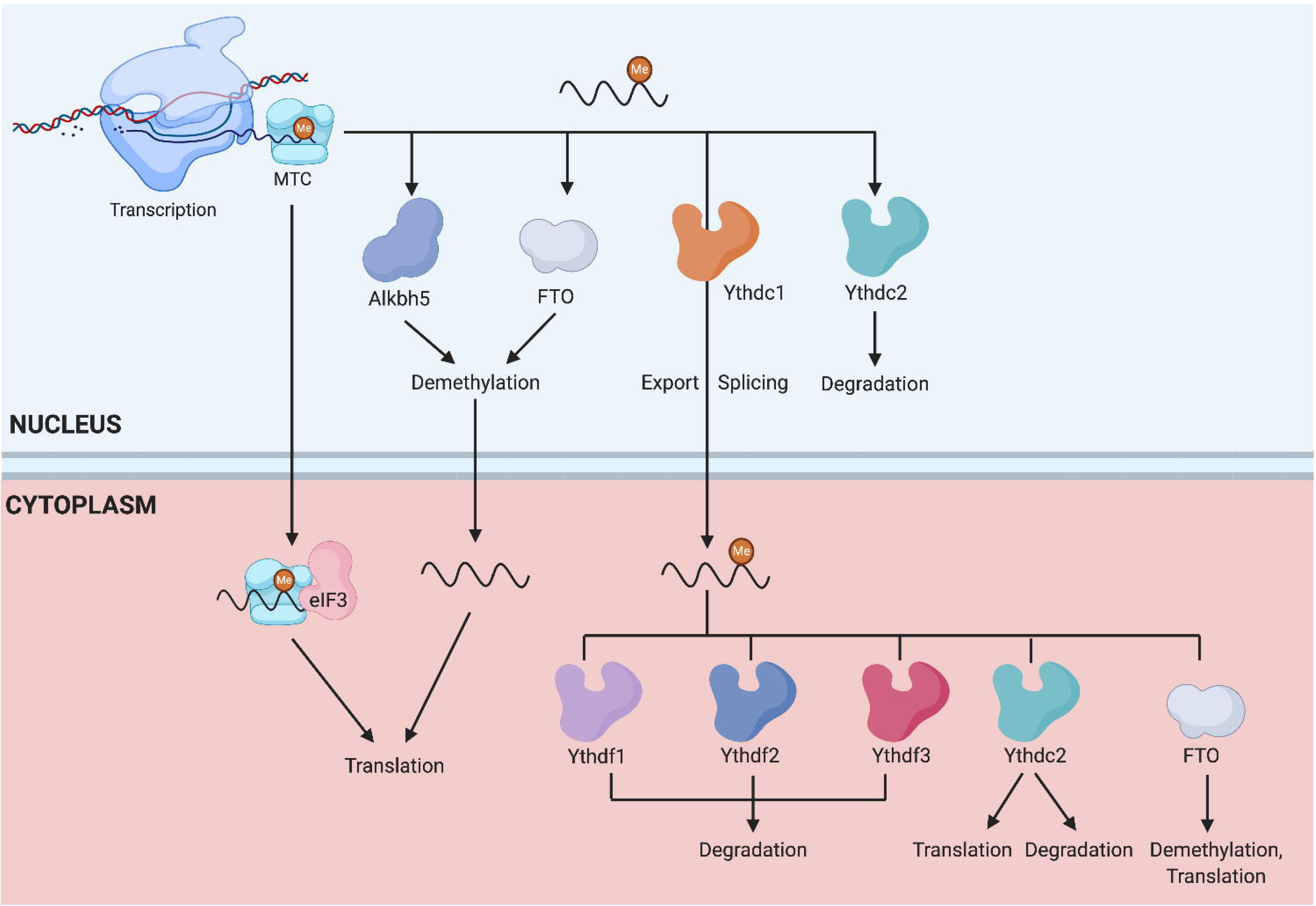
The regulation of m6A-RNAs. The ‘writer’ enzymes which form the methyltransferase complex (MTC) deposit methyl groups on the adenosine nucleotides of newly transcribed RNAs. These m6A-RNAs can remain complexed with the MTC and exported out into the cytoplasm for translation, demethylated by ‘eraser’ enzymes such as FTO and ALKBH5, undergo splicing and export by YTHDC1 or degradation by YTHDC2. In the cytoplasm, m6A-RNAs are recognized by ‘reader’ proteins (e.g. YTHDF1/2/3 and YTHDC2) which mediate their degradation or translation initiation. The modified RNAs can also undergo demethylation by cytoplasmic demethylase FTO. The compartmentalised expression of ‘writers, readers and erasers’ is likely to be dynamically regulated; the cell can distribute and redistribute these proteins between nuclear and cytoplasmic/synaptic compartments in a context-dependent manner.

### The m6A-Methylome as a Regulator of Intraneuronal Localisation

Gene expression in neurons is highly restricted by space and time ^71^. As such the subcellular pre-localisation of mRNAs to axons and dendrites not only saves energy but also ensures ‘on-demand’ translation coupled to activity-dependent signalling pathways, in a stimulus-, transcript-specific and spatially restricted manner ^72,73^. In this regard, m6A may travel with newly transcribed mRNAs or even direct their transport and serve as an epitranscriptomic-marks for local translation ^74^. Indeed, we have previously identified a composite of 4,469 m6A sites distributed to 2,921 genes across the pre- and post-synaptic compartments as well as the surround glial terminals ^17^. There were 1,266 hypermethylated synaptic genes which were enriched for a number of different functions such as synapse assembly, maturation, organisation, modulation of transmission, whereas the hypomethylated synaptic genes are enriched for metabolic functions. This functionally differential methylation for synaptic genes further supports a potential role of m6A in tagging transcripts for activity-dependent translation/decay that may be mediated by the expression of ‘readers’ ^17,33^.

Additionally, m6A machinery may also be localised to specific subregions. As mentioned above, Fto mRNA can be locally translated in axons in response to stimuli such as the nerve growth factor to demethylate Gap43 mRNA which leads to axonal elongation and axon growth ^75^. Similarly, axonal YTHDF1 has a role in facilitating the translation of m6A-mRNAs in crossing spinal commissural neurons and its deletion has been shown to result in the mis-projection of pre-crossing axons into motor columns ^76^.

Taken together, the accumulating evidence indicates an important role of m6A in maintaining the ‘regional autonomy’ and functionality of distal, non-nuclear compartments in the form of ‘ready-to-process’ pools of m6A-transcripts as well as the m6A machinery that are perhaps critical for energy-efficient, ‘on-demand’ responses to stimuli such as stress ^77^. The precise operational mechanisms via which transcripts are actually transported is unknown but could involve phase separation ^35–37^.

### Noncoding RNA interactions with the m6A-methylome

#### MicroRNAs

MicroRNAs (miRNAs) are small (21-23nt) non-coding RNAs but with a significant role in RNA silencing pathways including mRNA degradation, translational repression and transcriptional silencing ^78^. They are abundantly expressed in the brain where they have been implicated in a number of critical processes including neurodevelopment, synaptic plasticity as well as in the pathophysiology of major depression and PTSD ^79^. MiRNAs are derived from primary microRNAs (pri-miRNAs). Evidence has shown that m6A marks these pri-miRNAs for DGCR8-mediated cleavage into the mature miRNAs through m6A reader, HNRNPA2B1 ^80^. Depletion of METTL3 in vitro reduced the binding of DGCR8 to pri-miRNAs and resulted in a global reduction of mature miRNAs and concomitant accumulation of unprocessed pri-miRNAs ^81^. Even though mature microRNAs are short in length, recent evidence has shown that they too may harbour m6A in their sequence ^82^.

There is accumulating evidence for interactions between microRNAs and the m6A-methylome beginning with observations by Meyer et al. ^13^ who reported that 67% of 3’UTRs with m6A sites in mammalian RNAs contain at least one microRNA prediction target. Shortly after, Chen et al. reported that (a) deposition of m6A on target RNAs is dependent on microRNAs, (b) microRNAs are able to induce de novo m6A-methylation and (c) microRNA expression can influence nuclear localisation of METTL3 ^83^. Moreover, by virtue of its ability to regulate RNA splicing, m6A can also regulate the length of 3’UTRs and thus the availability of miRNA targets ^20^.

Transcripts for ‘writers’, ‘readers’ and ‘erasers’ are themselves targets of microRNAs. For example, miR-145 was found to negatively regulate the expression of YTHDF2 in hepatocellular carcinoma tissue ^84^. To further explore the potential interactions between microRNAs and the m6A machinery, we used the microRNA database miRNet ^85^ and identified 132 unique microRNAs which are able to regulate expression of METTL3, METTL14, YTHDC1, YTHDC2, YTHDF1, YTHDF2, YTHDF3, FTO and ALKBH5 in the human brain (Supplementary Table 1). As can be seen in the network plot (Figure 2), some of these microRNAs are presumably able to regulate expression of more than one of these targets which is consistent with their promiscuous mechanism of action.

**Figure 2.**
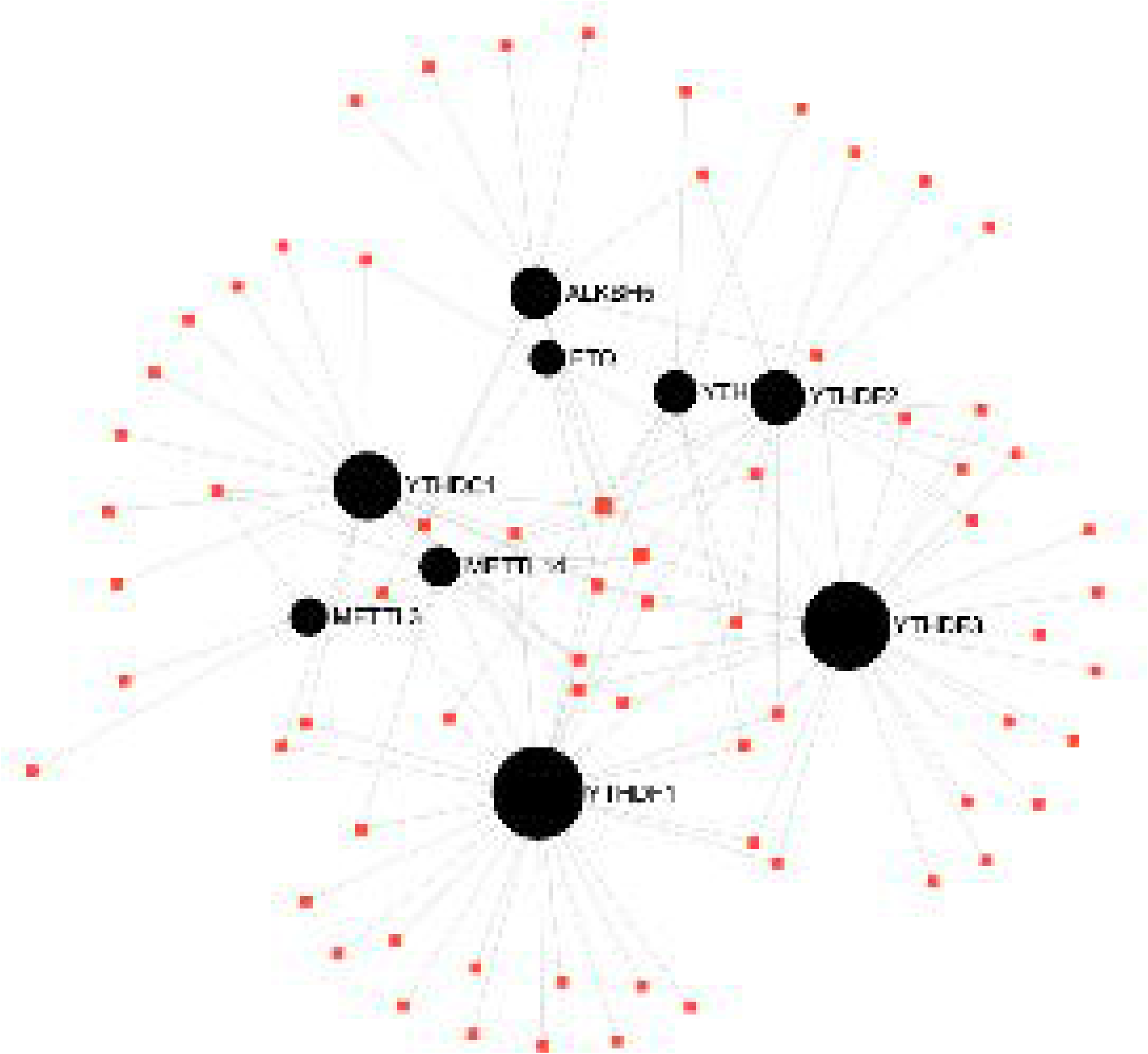
Network of the microRNAs (red squares) which regulate expression of m6A machinery genes (black circles). The size of the circles reflects the number of microRNAs which are predicted to regulate expression in humans. Generated using miRNET 2.0.

#### Long Non-Coding

Long non-coding RNAs (lncRNAs) are longer (>200nt) non-coding RNAs. Their synthesis profile has some features that are very similar to mRNAs, including splicing and the formation of secondary structures. Amongst their many functions as post-transcriptional regulators of gene expression, they can act as precursors to microRNAs, regulate their function by acting as decoys, compete with binding sites with mRNAs or by forming complexes with microRNAs ^86–88^. LncRNAs have been implicated in the pathophysiology of depression and PTSD ^89–91^.

The mechanistic evidence for interaction between lncRNAs and m6A machinery is scarce with only study so far showing that the lncRNA FOXM1-AS facilitates the interaction between FOXM1 mRNA and ALKBH5 which leads to increased expression of FOXM1 ^92^. However, there are many more studies which have reported on the impact of epitranscriptomic modifications including m6A on lncRNA function ^93^. For example, m6A-methylation of the lncRNA metastasis associated lung adenocarcinoma transcript 1 (MALAT1) was found to influence the physical access (see above ‘m6A-structural switch) to RNA-binding proteins ^94^; m6A-methylation of the lncRNA X-inactive specific transcript (XIST) and its subsequent recognition by YTHDC1 was a precursor to transcriptional silencing ^22^. While long intergenic non-coding RNAs (lincRNAs) are also known to harbour m6A ^95^, evidence for the functional role of m6A in lincRNAs is limited except for one study which showed that m6A-methylation of linc1281 was necessary for differentiation of mouse embryonic stem cells ^96^.

### Stress-induced Psychopathologies and the m6A-Methylome

The neuroendocrine response to stress, also known as the HPA axis, involves the initial release of corticotropin releasing factor (CRF) by the paraventricular nucleus of the hypothalamus and amygdala, which stimulates CRF receptors (CRFR) on the anterior pituitary to release adrenocorticotropic hormone (ACTH), ultimately resulting in the release of corticosteroids by the adrenal cortex. Corticosteroids can bind to gluco-(GR) and mineralocorticoid receptors (MR) on a range of target organs including the brain to initiate fast non-genomic effects (e.g. glutamate release, endocannabinoid release) as well as slower genomics effects ^97,98^.

In this section, we have firstly reviewed evidence from animal studies which demonstrate the impact of stress on the m6A-methylome/machinery and its relationship to stress-induced behavioural changes of relevance to major depression and PTSD (Table 2). Stressors can be either acute or chronic but the difference in impact is rather diffuse ^98^. For the discussion below on major depression, we have focused on models which utilise restraint stress or chronic unpredictable stress. For the discussion on PTSD, we have chosen models which use variations of the fear conditioning paradigm. We then examined human studies which report evidence for a dysregulated m6A-methyome/machinery in these two disorders, incorporating a bioinformatic meta-analysis of publicly available datasets (Supplementary Table 2).

**Table 2.**
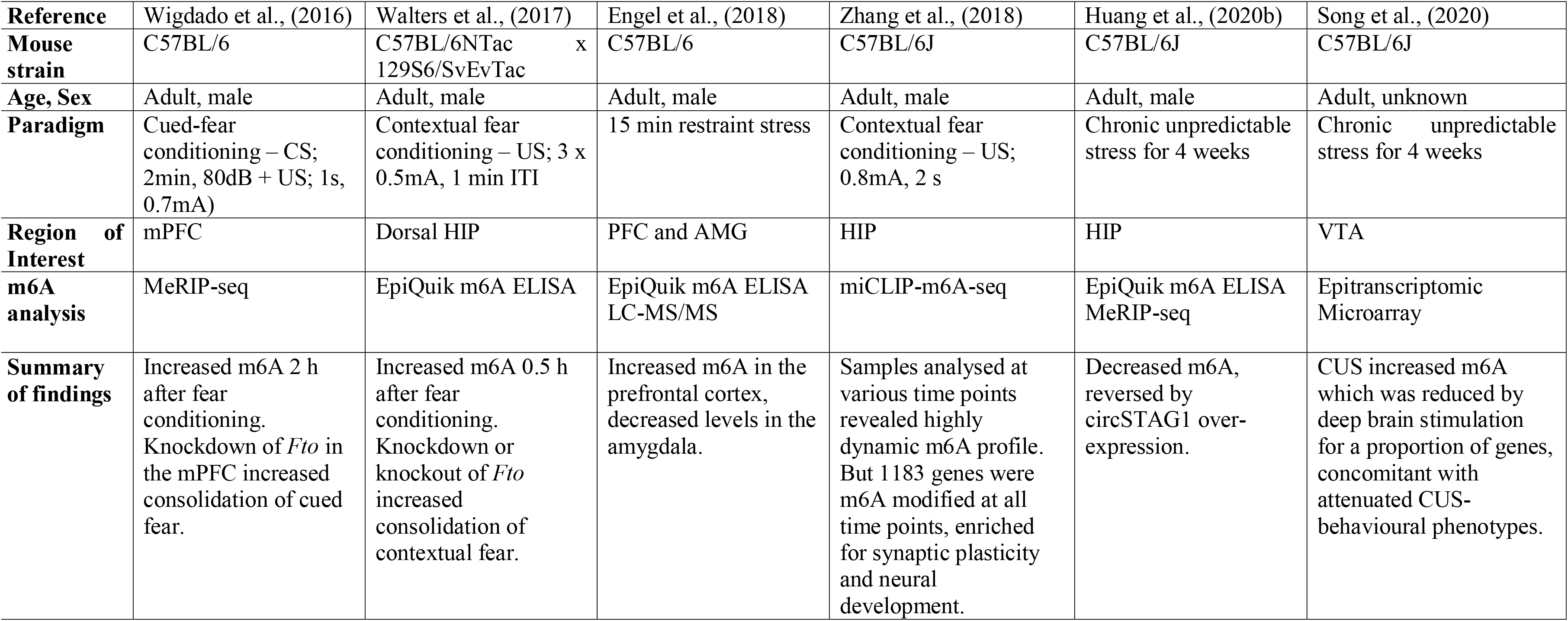
Summary of preclinical studies which have used stress paradigms to assess impact on m6A profile in various brain regions.

|  |  |  |  |  |  |  |
| --- | --- | --- | --- | --- | --- | --- |
| **Reference** | Wigdado et al., (2016) | Walters et al., (2017) | Engel et al., (2018) | Zhang et al., (2018) | Huang et al., (2020b) | Song et al., (2020) |
| **Mouse strain** | C57BL/6 | C57BL/6NTac x 129S6/SvEvTac | C57BL/6 | C57BL/6J | C57BL/6J | C57BL/6J |
| **Age, Sex** | Adult, male | Adult, male | Adult, male | Adult, male | Adult, male | Adult, unknown |
| **Paradigm** | Cued-fear conditioning – CS; 2min, 80dB + US; 1s, 0.7mA) | Contextual fear conditioning – US; 3 x 0.5mA, 1 min ITI | 15 min restraint stress | Contextual fear conditioning – US; 0.8mA, 2 s | Chronic unpredictable stress for 4 weeks | Chronic unpredictable stress for 4 weeks |
| **Region of Interest** | mPFC | Dorsal HIP | PFC and AMG | HIP | HIP | VTA |
| **m6A analysis** | MeRIP-seq | EpiQuik m6A ELISA | EpiQuik m6A ELISA LC-MS/MS | miCLIP-m6A-seq | EpiQuik m6A ELISA MeRIP-seq | Epitranscriptomic Microarray |
| **Summary of findings** | Increased m6A 2 h after fear conditioning. Knockdown of *Fto* in the mPFC increased consolidation of cued fear. | Increased m6A 0.5 h after fear conditioning. Knockdown or knockout of *Fto* increased consolidation of contextual fear. | Increased m6A in the prefrontal cortex, decreased levels in the amygdala. | Samples analysed at various time points revealed highly dynamic m6A profile. But 1183 genes were m6A modified at all time points, enriched for synaptic plasticity and neural development. | Decreased m6A, reversed by circSTAG1 over-expression. | CUS increased m6A which was reduced by deep brain stimulation for a proportion of genes, concomitant with attenuated CUS-behavioural phenotypes. |

#### Major Depression – Animal Studies

In rodents, many strategies exist to induce aspects of depression, which includes exposure to a restraint or chronic unpredictable stress (CUS) paradigm. Engel et al. ^28^ analysed epitranscriptomic changes 4 h after 15 mins of restraint stress with mouse cortical RNA samples using m6A-seq. They showed at baseline that half of all expressed cortical genes were m6A methylated and most of these were implicated in synaptic plasticity and neuronal regulation. However, 4 h after restraint stress, the magnitude of differential gene expression compared to controls was low and furthermore, the impact on cortical m6A levels was also insignificant; the authors speculate that these findings were likely to be due to heterogenous nature of input material which may have precluded the identification of regional effects of the restraint stress. Based on these results, the authors focused on examining the epitranscriptome on total RNA from the prefrontal cortex (PFC) and the amygdala (AMG). At 30 mins after the stress, there were no differences in global m6A profiles. But from 1 h to 24 h the m6A levels were higher in the AMG and lower in the PFC; these results were confirmed for up to 4 hrs using isolated mRNA and LC-MS. This tissue-specific result was matched by lower mRNA expression of *Fto*, *Alkbh5* and *Ythdc1* in the AMG than in the PFC over the same time period. The impact of acute restraint stress on global m6A/m levels was also detected in the blood along with altered mRNA expression of *Mettl3*, *Wtap* and *Alkbh5*. The authors then applied m6A-RIP-qPCR to show differences in the absolute methylation of a number of specific stress and synaptic plasticity-related transcripts in the PFC and AMG. Interestingly, they reported a negative correlation between stress-induced changes in gene expression and absolute m6A/m levels which led to the suggestion that these modifications were mainly linked to transcript decay ^28^.

Huang et al., ^68^ reported a CUS-induced reduction in hippocampal m6A levels and m6A-seq revealed a greater down-regulation in m6A-methylation than up-regulation of a number of transcripts, including for fatty acid amide hydrolase (*Faah)* mRNA. CUS also induced an increase in hippocampal (HIP) FAAH protein expression. This enzyme is a major component of the endocannabinoid system which metabolises the archetypal endocannabinoid, anandamide (AEA) ^99^ and is also critical to the stress response ^100^. Sustained FAAH activity have been associated with maladaptive stress responses ^101^. The authors also reported that there was CUS-induced localisation of ALKBH5 in the nucleus of astrocytes whereas under control conditions, ALKBH5 is normally bound to the circular RNA, *circSTAG1*, in the cytoplasm. The expression of *circSTAG1* in the HIP, blood and plasma of CUS mice was lower than controls and this profile was matched by similar differential expression in the blood and plasma in patients with major depression. Overexpression of *circSTAG1* captured and restricted ALKBH5 to the cytoplasm, preventing the demethylation of nascent *Faah* mRNA molecules, degradation in the cytoplasm and the attenuation of depressive-like behaviours. It is worth highlighting that other transcripts of interest with significantly decreased m6A-methylation include *Nape-pld*, an enzyme involved in the synthesis of endocannabinoids, *Fkbp5* as well as *Alkbh5* ^68^.

Another recent study also used the CUS paradigm in mice to examine the impact on the m6A-methylome in the ventral tegmental area (VTA), another stress-sensitive brain structure involved in the reward system ^102^. In contrast with the previous study, CUS-induced a significant increase in m6A-methylation of transcripts in the VTA. Deep brain stimulation (DBS), which is reportedly an effective form of treatment for major depression, reversed CUS-induced depressive-like behaviours as well as changes in m6A-methylation status for 329 genes. Notably, there were also a number of genes for which the expression was not altered between CUS and CUS-DBS treated mice, but the m6A-methylation status was significantly different. A correlation analysis also revealed no significant relationship between m6A-methylation status and gene expression.

#### Major Depression – Human Studies

To date, there has been only one study to quantify m6A levels in patients suffering from major depression. Engel et al., ^28^ first administered oral dexamethasone to healthy controls and observed a reduction in m6A/m levels, mRNA expression of *METTL3*, *FTO*, *ALKBH5*, *YTHDF1*, *YTHDF2* and an increase in *YTHDF3* in the blood. They then administered a dexamethasone challenge test to healthy controls and patients, as a way of assessing glucocorticoid receptor resistance ^103^, and observed a reduction in global m6A/m in blood only in the controls. These results were confirmed by similarly treating B lymphocyte cell lines (BLCLs) derived from controls and patients with dexamethasone or cortisol. They then performed m6A-seq on control and patient BLCLs treated with cortisol, observing that there were more cortisol-regulated m6A peaks in BLCLs from controls than patients. Additionally, m6A-RIP-qPCR revealed a significant decrease in absolute m6A/m for several stress response genes, including *FKBP5*, following cortisol treatment in BCLs from controls but not patients. FKBP5 is a chaperone protein which modulates activity of the glucocorticoid receptor and has been implicated in stress-induced psychopathologies ^104^. In the absence of differences in expression of the glucocorticoid receptor in the blood, these results led to the conclusion that signalling processes downstream of receptor activation are likely to be altered in patients, affecting the methylation status of stress response transcripts such as *FKBP5* ^28^.

A number of studies have identified single nucleotide polymorphisms (SNP) in the *FTO* gene which appears to moderate a complex relationship between obesity and major depression ^105–108^. At least one of these SNPs (rs9939609) has a limited association with major depression^109^. A SNP in *ALKBH5* gene has also been associated with depression ^110^.

To further explore the potential role for a dysregulated m6A-methylome in major depression, we identified and analysed relevant publicly available single-nucleus (sn) and bulk RNA-seq datasets from the NCBI GEO database (see Supplementary Info for Methods). We hypothesized that the features of the pathophysiology of depression may include altered expression of m6a ‘writers, readers and erasers’.

The snRNA-seq study consisted of gene expression profiles of ~70000 cells derived from prefrontal cortex (PFC) of post-mortem brain samples of 17 patients diagnosed with major depression who had died due to suicide and 17 matched healthy controls who died due to accidents or natural causes ^111^. Heatmap of mean log normalised expression for m6A machinery genes in different PFC cell subtypes is represented in Figure 3. Genes encoding ‘writer’ *METTL16*, ‘readers’ *YTHDC1* and *YTHDC2* (marked by yellow boxes), show significant differential expression in three neuronal cell types. Specifically, *METTL16* was upregulated in both L5/6 excitatory pyramidal neurons and inhibitory somatostatin neurons in patients. *YTHDC1* and *YTHDC2* were down- and upregulated respectively, in inhibitory somatostatin neurons.

**Figure 3.**
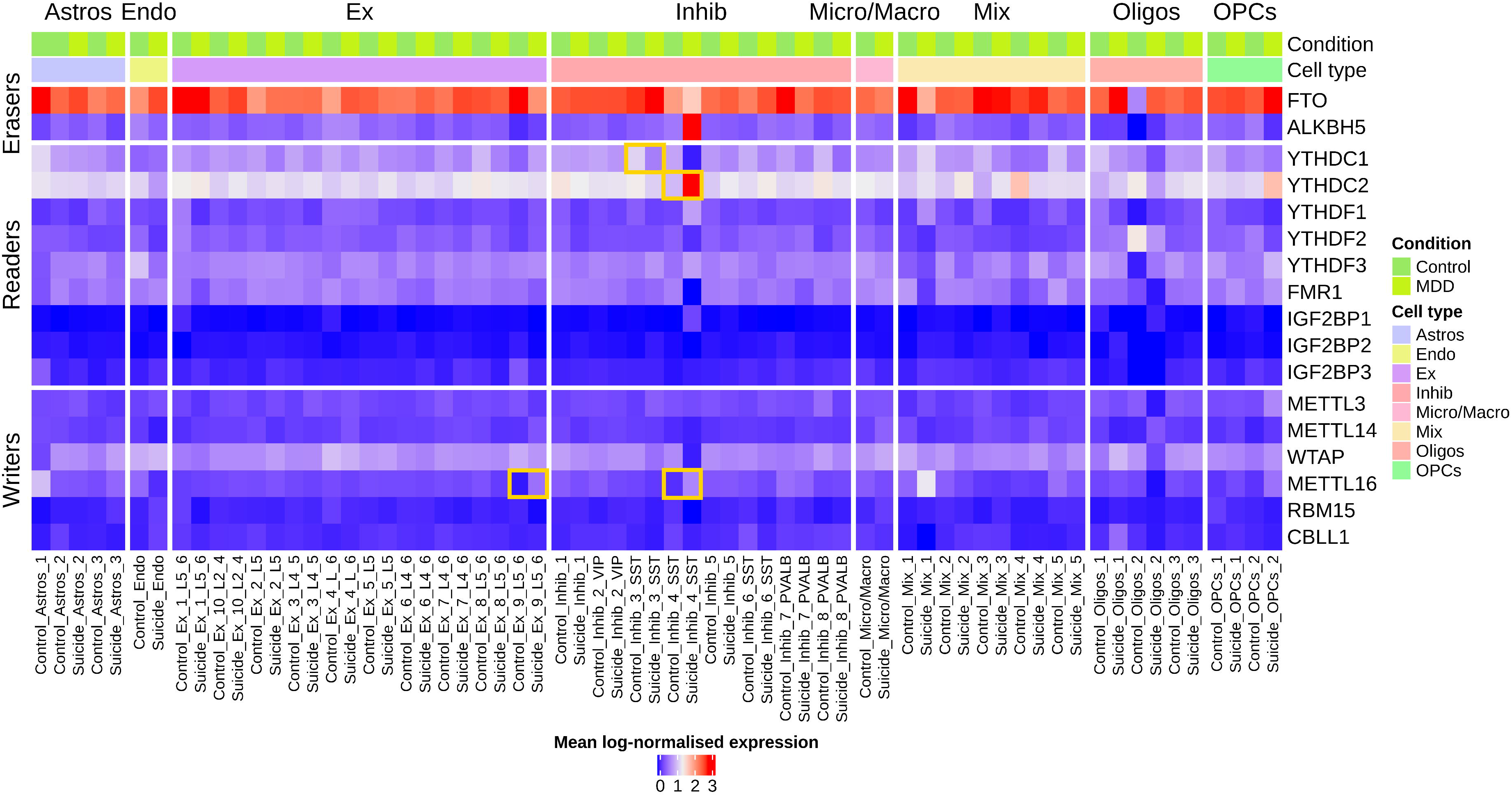
Heatmap of snRNA-seq mean log-normalised expression for m6A machinery genes in the PFC of major depression patients and healthy controls. The four yellow boxes highlight differentially expressed m6A ‘readers’, YTHDC1/2 and the m6A ‘writer’ METTL16 in somatostatin inhibitory neurons and excitatory neurons (p<0.05). Astros: astrocytes; Endo: endothelia; Ex: Excitatory neurons; Inhib: Inhibitory neurons; Micro/Macro: Microglia/Macrophage; Oligos: Oligodendrocytes; OPCs: Oligodendrocyte precursor cells.

Bulk transcriptomics datasets from several other brain regions and noted significant differential expression of ‘writers,’ ‘readers,’ and ‘erasers’ was detected in examined structures except for PFC (Figure 4).

**Figure 4.**
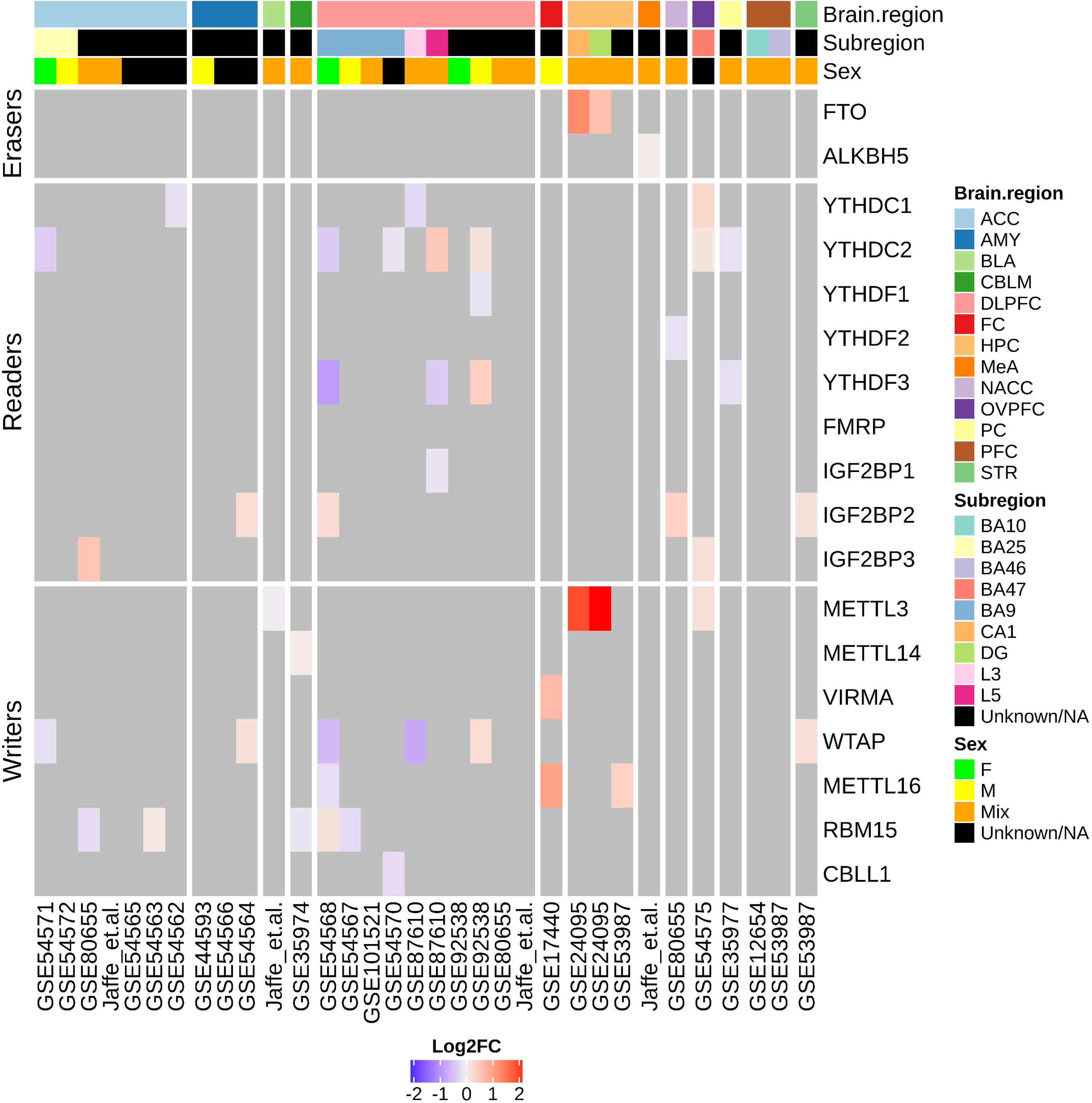
Heatmap of log2FC for m6A machinery gene expression in bulk RNA-seq/microarray expression profiling experiments in patient samples vs controls from different brain regions and conditions. Top three index bars describe the brain regions, subregions, sex of the subjects and the bottom row indicates the reference or NCBI GEO series ID. Black bars indicate either no subregion analysis or unspecified details on the sex of the subjects in the relevant studies. Grey areas in the heatmap indicate no differential gene expression. F: females; M: males; Mix: both) (ACC: anterior cingulate cortex; AMY: amygdala; CBLM: cerebellum; DLPFC: dorsolateral prefrontal cortex; FC: frontal cortex; HPC: hippocampus; NACC: nucleus accumbens; OVPFC: orbital ventral prefrontal cortex; PC: parietal cortex; PFC: prefrontal cortex; STR: striatum.

Given the influence of the m6A-methylome on gene expression, we next investigated if there was a relationship between the number of m6A sites per gene and differential gene expression in major depression. For this, we used an m6A-seq dataset generated from human cerebrum and cerebellum and exomePeak to determine the number of m6A sites in each gene ^3,112^. Using the bulk transcriptomics datasets above, we then grouped differentially expressed genes in patients vs controls based on the m6A peaks count in respective brain regions. Log2 fold change (log2FC) values (patients vs controls) for genes were presented in cumulative distribution function plots (Figure 5). The analysis showed sex-specific effects as well as effects between and within brain regions. These have been briefly discussed below. The findings from other analyses have been discussed in the Supplementary Information (Supplementary Figure 1).

**Figure 5.**
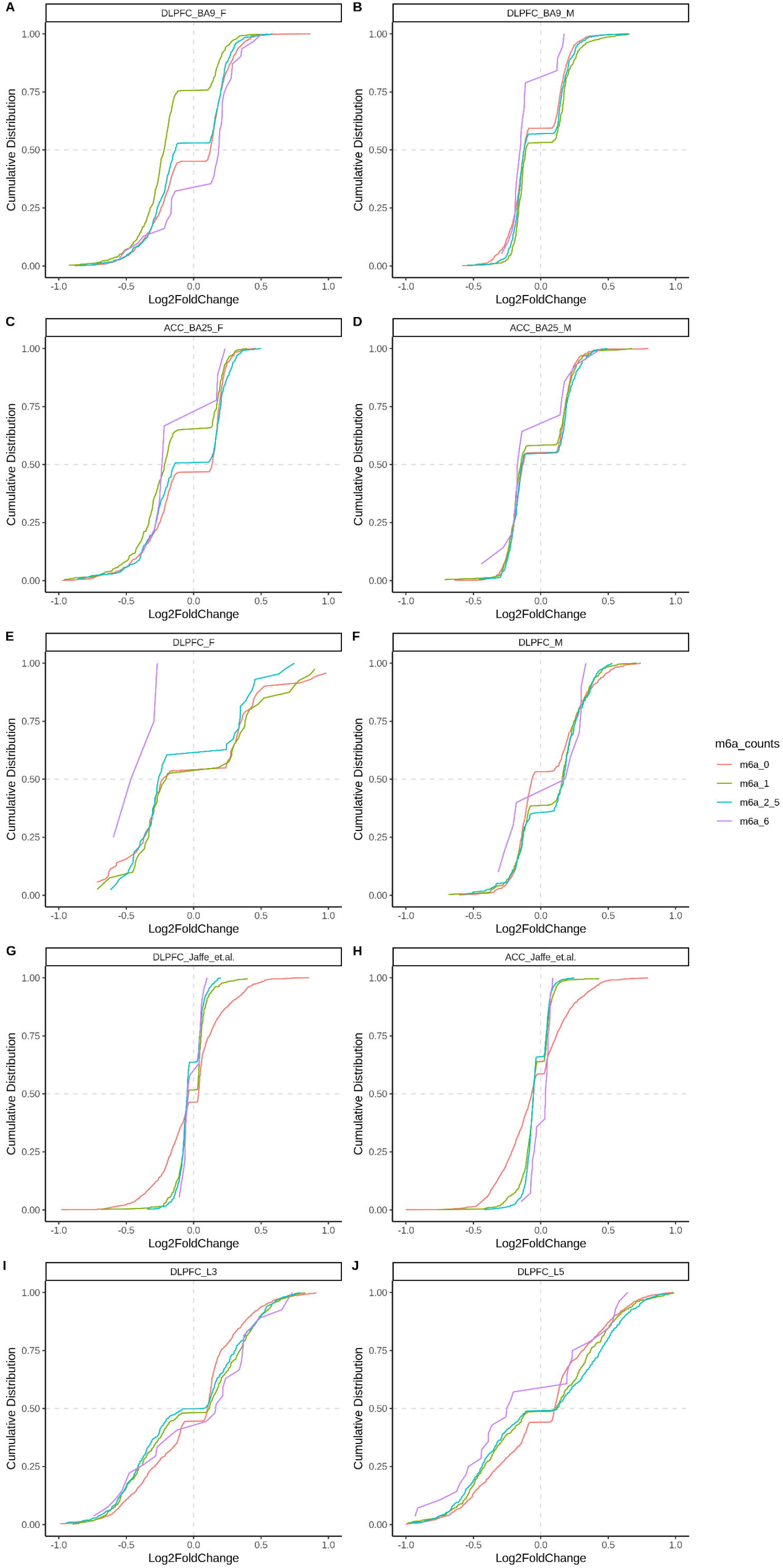
Cumulative distributions of log2FC of gene expression in MDD patients vs control show sex- and brain region-dependent regulation. The differentially expressed genes are divided into four groups based on the number of m6a sites mapped to their RNA. m6a_0: no m6a peaks; m6a_1: one m6a peak; m6a_2_5: two to five m6a peaks; m6a_6: six or more m6a peaks. DLPFC: dorsolateral prefrontal cortex; ACC: anterior cingulate cortex.

The relationship between differential gene expression and the number of m6A sites per gene was highly dynamic in females compared to males across and within brain regions (Figure 5A-F). For example, in the study by Chang et al., ^113^, in the BA9 region of dorsolateral prefrontal cortex (DLPFC) female samples, there was a larger distribution of genes with one m6A site that were downregulated. Similarly in the BA25 region of anterior cingulate cortex (ACC) female samples, there was a larger distribution of genes with more than five m6A sites which showed downregulation as represented by the higher cumulative distribution of genes for log2FC lower than zero (Figure 5A, C). However, in matched male samples for the BA9 region of DLPFC, there was a larger distribution of genes with more than five m6A sites that were downregulated as compared to remaining genes; this was also the case in the BA25 region of ACC for males (Figure 5B, D). In another study which also analysed the DLPFC of males and females, but with no layer information, there was a larger distribution of downregulated genes with more than five m6A sites in females (Figure 5E) ^114^. However in males, there was a uniform distribution of log2FC values for genes with more than five m6A sites (Figure 5F).

In comparing the DLPFC and ACC, the differences in differential gene expression for genes with five or less m6A sites was comparable. However, for genes with more than five m6A sites, there was a greater proportion of upregulated genes in the ACC than in the DLPFC (Figure 5G-H) ^115^. In comparing gene expression in L3 and L5 pyramidal neurons of DLPFC, our analysis of a third study showed that genes containing m6A sites had similar distribution of log2FC in L3 pyramidal neurons but in L5 pyramidal neurons there was a greater distribution of genes with more than five m6A sites that were downregulated compared to remaining groups (Figure 5I-J) ^116^.

#### Post-Traumatic Stress Disorder – Animal Studies

The psychology of PTSD is rooted in the acquisition of experiences and formation of memories associated with specific traumatic events. Accordingly, persistent changes in brain structures such as the HIP and AMG with roles in learning and memory have been implicated in the pathophysiology of this disorder ^117^. A commonly used paradigm to induce aspects of PTSD in rodents involves fear conditioning. It must be noted however that the focus of the majority of studies reviewed below was to examine the role of the epitranscriptome in the context of learning and memory with unconditioned stimuli (US) that are weaker in intensity compared to those used to induce long-lasting, robust phenotypes of relevance to PTSD in rodents ^118^. This caveat imposes limits on the conclusions that can be drawn from the studies in the context of this disorder.

In an early study by Widagdo et al., ^119^, a cued-fear conditioning procedure was used in which adult male mice were exposed to an auditory conditioned stimulus (CS, tone) paired with a US (0.7 mA, 1 s). Each mouse received 6 x CS-US pairs with an intertrial interval of 2 min. The mice were sacrificed 2 h later and RNA from the medial prefrontal cortex (mPFC) was analysed with m6A-seq. Results showed that compared to naïve mice, there was a significant increase in the number of m6A peaks in the CS-US mice for mPFC genes involved in synaptic plasticity and most were located within the vicinity of the stop codon. The fear conditioning paradigm caused no change in the mRNA expression of *Mettl3* or *Mettl14* but decrease in *Fto*. The authors then used a cortical neuronal culture system and m6A-IP-qPCR to validate the increased m6A levels for several of the identified plasticity-related genes. Using this in vitro setup, they also showed that the over-expression or knocking down of FTO was able to alter the decay rate of specific transcripts. And in their final experiment, they knocked down *Fto* in the mPFC of mice and showed that while this manipulation had no impact on fear acquisition, it did affect memory consolidation with the mice showing increased freezing behaviour in response to the cue 24 h after the training ^119^.

Another early study by Walters et al. ^120^ used a contextual fear conditioning paradigm. Mice were placed inside the fear conditioning chamber and exposed to three 0.5 mA foot shocks with 1 min intervals between shocks. The mice were sacrificed 0.5 or 1 h later, and the HIP was used for analysis. At the 0.5 h time point, there was no effect in m6A levels in total RNA but a significant increase in m6A levels in mRNA. There was also a significant decrease in expression of FTO, mainly in the synaptosome (a tissue preparation with enriched synaptic terminals). The authors then used CRISPR-Cas9 and a knockdown approach targeting *Fto* in the dorsal HIP, to demonstrate that it was involved in the consolidation of contextual fear memories ^120^.

Following on from their examination of the impact of acute restraint stress on the m6A-methylone, Engel et al., ^28^ generated *Mettl3* and *Fto* conditional knockout mice deficient in expression in excitatory neurons in the neocortex and HIP. They reported that while absence of Mettl3 caused a reduction in global m6A levels, conditional deletion of *Fto* had no effect. They also generated conditional knockout mice with a timed deletion of *Mettl3* and *Fto* in pyramidal neurons of the dorsal and ventral HIP. The baseline behavioural phenotypes of these mice were comparable to wildtypes, except for increased and decreased marble burying behaviour in the *Mettl3* and *Fto* conditional knockout mice, respectively. All conditional knockout mice and wildtypes were subjected to a fear conditioning procedure with a single CS (80 dB tone, 20 s) paired with a US (0.7 mA, 2 s) and then assessed for cued-or contextual memories. There was no difference in acquisition compared to wildtypes but both conditional knockout mice displayed increased cued- and contextual fear memory that was resistant to extinction. This phenotype was underpinned by fear-conditioning induced effects on expression of genes for neurotransmission and transcription which were more pronounced in *Fto* conditional knockout mice than in the *Mettl3* conditional knockout mice compared to unstressed conditional knockout mice ^28^.

Using a contextual fear conditioning paradigm, Zhang et al. ^29^ showed that the number of m6A-modified genes in the HIP were dynamically regulated at 0.5, 1 and 4 h following contextual fear conditioning with a single US (0.8 mA, 2 s). However, there were 1183 genes that were m6A-modified at all time points and these were enriched for synaptic plasticity and neural development. These authors also showed that *Mettl3* conditional knockout mice displayed reduced freezing to the context 24 h but not 30 min after training which suggested a deficit in long-term memory formation. This deficit could be corrected by restoring METTL3 expression in the dorsal HIP. However, when these conditional knockout mice were exposed to three foot shocks instead of one, contextual fear memory was no different to wildtypes. Interestingly, in comparing the temporal effects of the fear conditioning (one shock) between wildtype and *Mettl3* cKO mice, there were no differentially expressed genes in the hippocampus over 4 hours. However, the absence of METTL3 led to a significant reduction in the number of m6A-modified transcripts for several immediate early genes. Further analysis revealed a significant reduction in the protein expression for these IEGs which led to the conclusion that METTL3 is necessary to modify and shunt m6A-IEG transcripts down a translational pathway to facilitate memory consolidation in the HIP following fear conditioning ^29^.

To investigate the functional roles of the m6A reader YTHDF1 in learning and memory, Shi et al., ^47^ generated CRISPR-Cas9 knockout mice. For fear conditioning, mice received either one of three different stimuli: a single pairing of a CS tone (75 dB, 30 s) with a US foot shock (0.5 mA, 1 s) for the weak protocol, a single pairing of the CS tone with a longer US (2 s) for the moderate protocol or three pairings of the CS tone with the longer US and an inter-trial interval of 1 min for the strong protocol. During the inter-trial interval with the moderate training protocol, *Ythdf1* knockout mice froze less than the wildtypes but not when the tone was presented, suggesting that contextual learning was impaired in these mice. *Ytdhf1* knockout mice displayed deficits in contextual fear memory when assessed 24 h but not 2 h later and these deficits were rescued with the hippocampal re-expression of YTDHF1. Contextual-fear memory was also reduced in these mice 2 h after the weaker training protocol but there were no genotype differences with the stronger protocol at 24 h ^47^.

#### Post-Traumatic Stress Disorder – Human Studies

To date, there have been no studies which have specifically profiled the m6A-epitranscriptome or expression of m6A machinery in patients with PTSD but similar to our approach above, we searched for and bioinformatically analysed publicly available datasets. We observed differential expression of m6A machinery genes in peripheral blood samples from PTSD patients compared to controls (Figure 6). There was a significant dysregulation of several ‘writers’ (GSE81761, GSE67663) and ‘readers’ (GSE81761, GSE860) but no effect on ‘erasers.’ Sleep treatment reduced expression of the ‘writer’ *RBM15* and the indirect ‘reader’ *IGF2BP2* (PTSD_Notimproved_vs_Improved, GSE81761) in patients who did not reported improvements in symptomatology. In patients who had been admitted to the emergency room (PTSD_vs_NoPTSD_ER) following trauma, there was a downregulation in *YTHDC1* and *YTHDF2* in peripheral blood mononuclear cells (PMBCs), but this difference was not detected 4 months later (PTSD_vs_NoPTSD_M4, GSE860).

**Figure 6.**
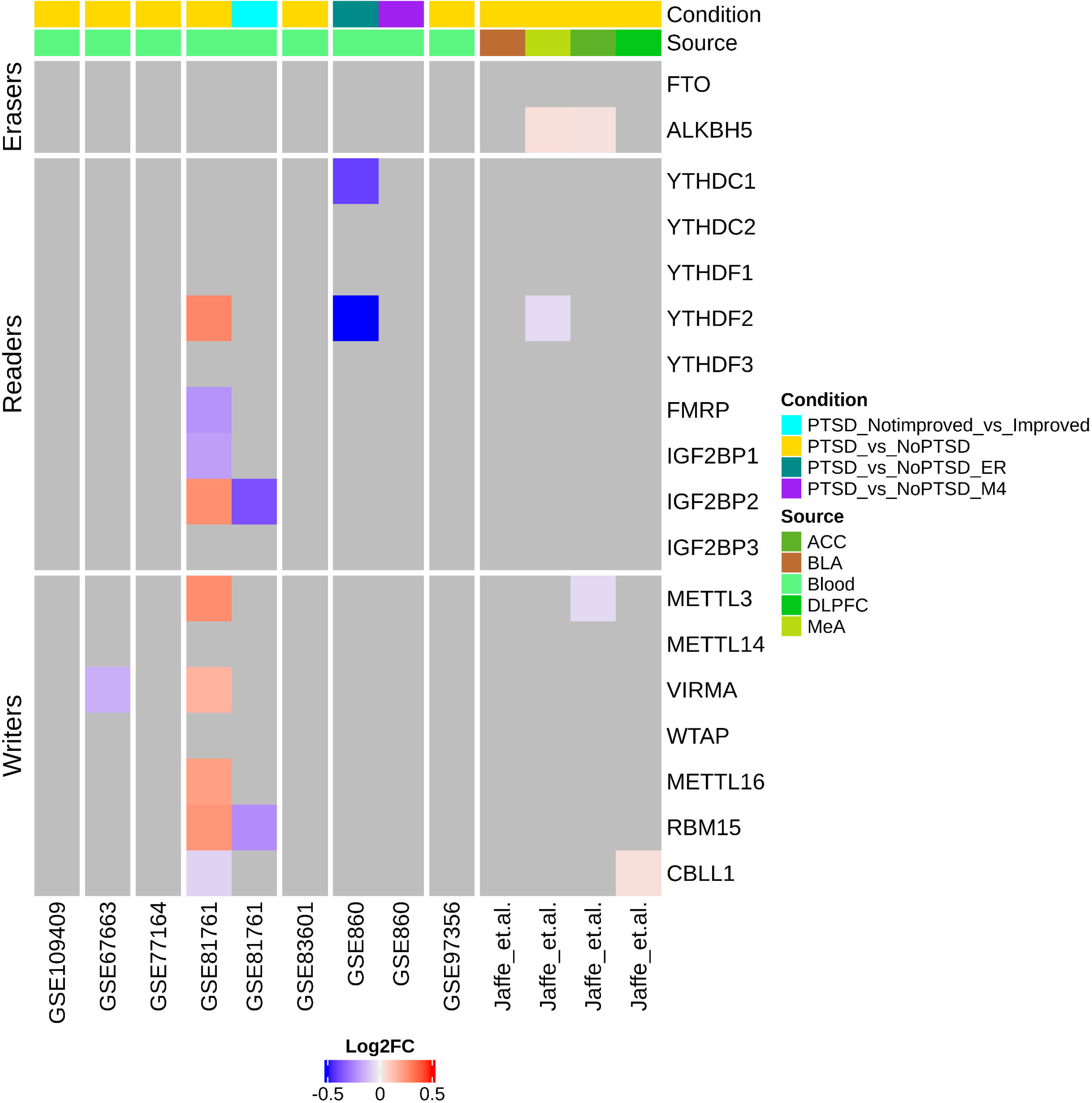
Heatmap of log2FC for m6A machinery gene expression in blood and brain regions of PTSD patients. Index bars above the heatmap indicates comparisons and sample source. BLA: basolateral amygdala; MeA: medial amygdala; ACC: anterior cingulate cortex; DLPFC: dorsolateral prefrontal cortex.

In a recent preprint, Jaffe et.al. ^115^ reported differentially expressed genes in PTSD patients as compared to controls after correcting for various factors in a cohort of 107 PTSD patients. We examined differential expression of m6A machinery genes in PTSD vs controls from four brain regions (viz. two cortical regions; DLPFC and ACC and two amygdala regions; basolateral amygdala (BLA) and medial amygdala (MeA)). Expression of the m6A ‘eraser’ ALKBH5 was significantly upregulated in ACC and MeA regions of the brain in PTSD patients as compared to control. Similarly, we also observed that components of the MTC such as *METTL3* and *CBLL1* were downregulated in the ACC and upregulated in the DLPFC, respectively (Figure 6). We constructed cumulative distribution function plots for these brain regions and observed that in the DLPFC, there was a larger distribution of upregulated genes with at least one m6A site compared to genes with no m6A sites which showed a uniform distribution of up and downregulated genes. In the ACC, there was a larger distribution of downregulated genes with no m6A sites and larger distribution of upregulated genes with two or more m6A sites (Figure 7).

**Figure 7.**
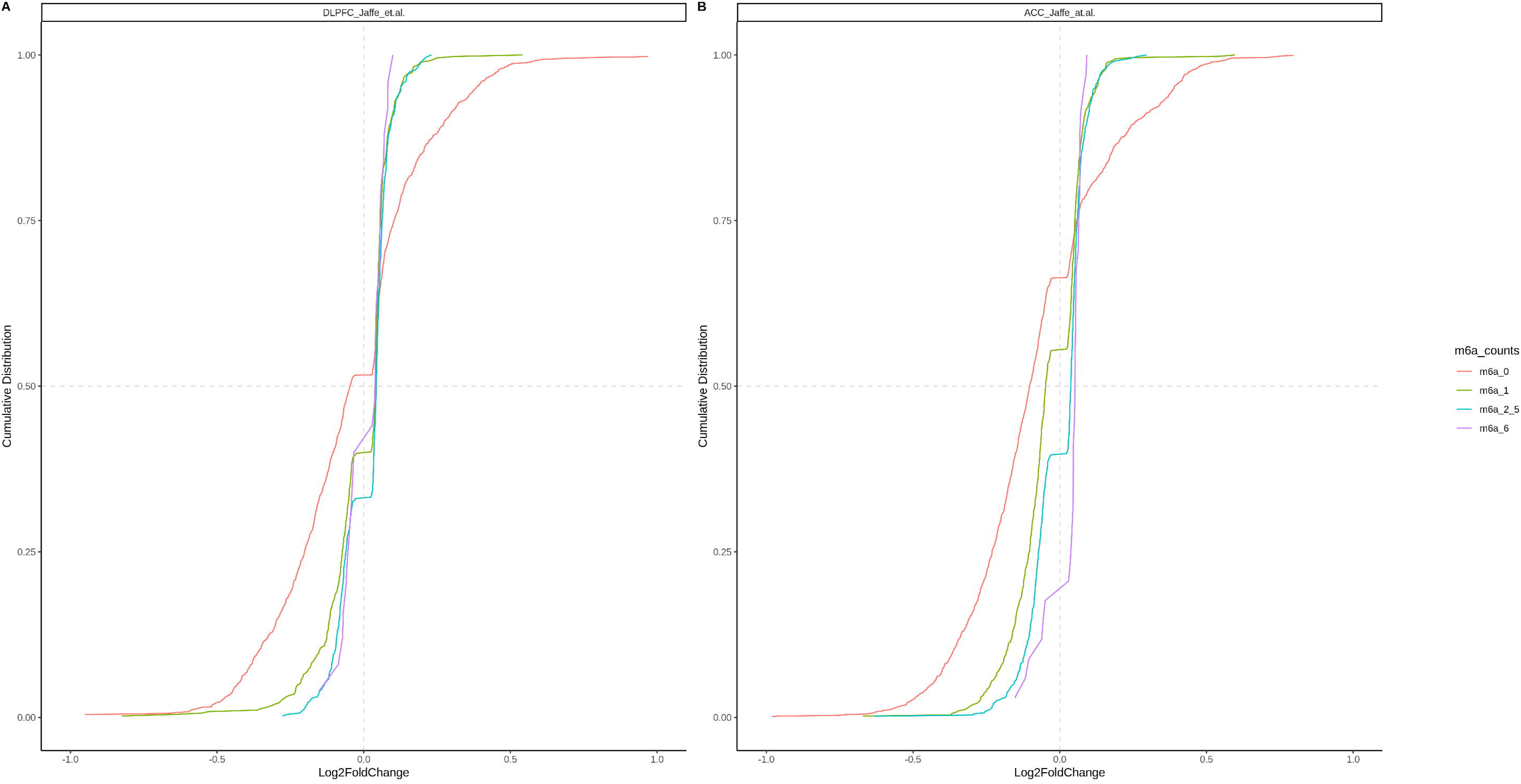
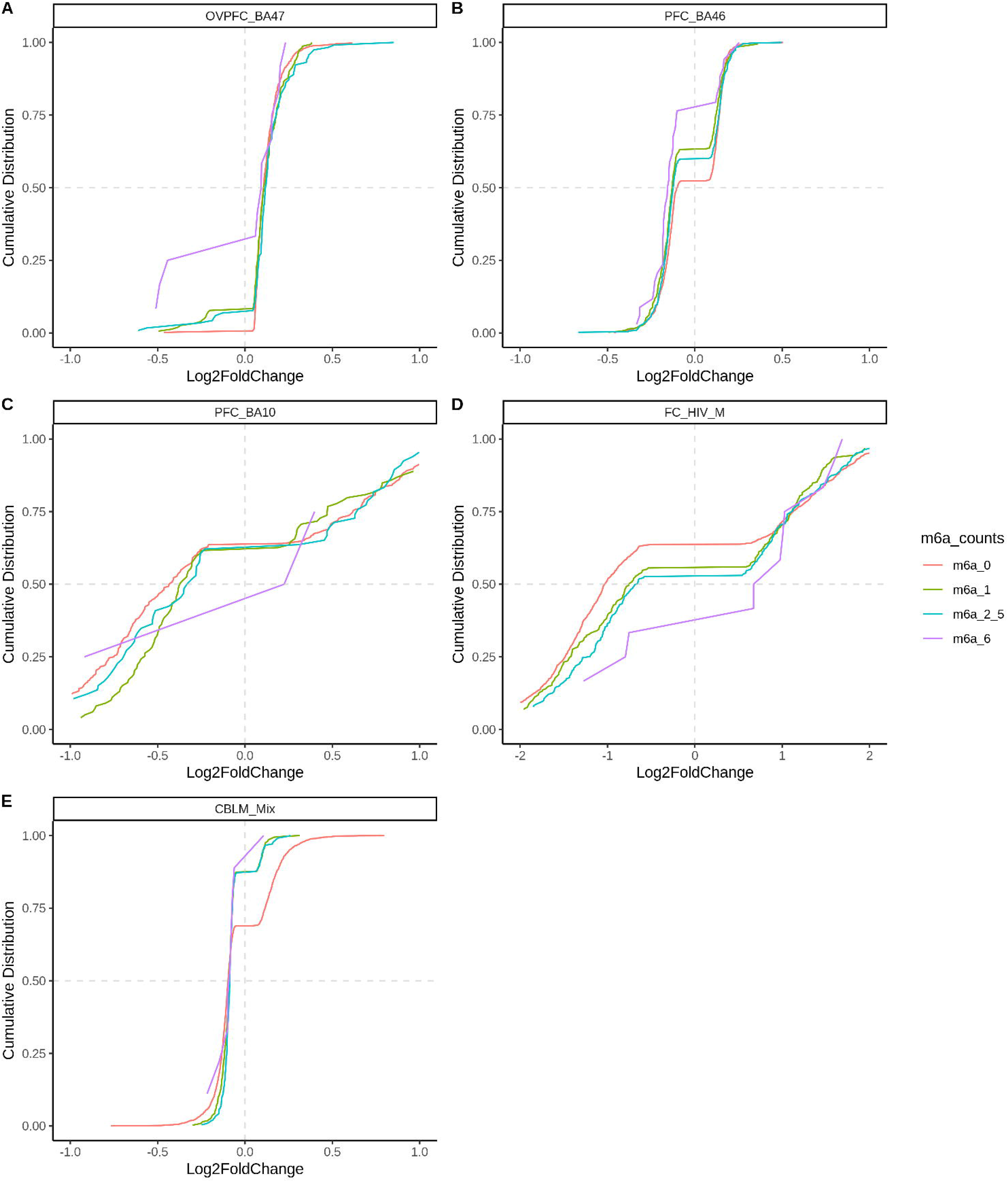
Cumulative distribution plot for log2FC of significant differential genes in PTSD patients as compared to controls in different conditions. The genes are divided in four groups based on the number of m6a sites mapped to their RNA. m6a_0: no m6a peaks; m6a_1: one m6a peak; m6a_2_5: two to five m6a peaks; m6a_6: six or more m6a peaks. DLPFC: dorsolateral prefrontal cortex; ACC: anterior cingulate cortex.

Overall, the evidence from animal model investigations strongly implicate a role for m6A-methylome/machinery genes in mediating aspects of the response to acute and chronic stress. Moreover, our bioinformatic analysis of publicly available human data for these disorders indicate that not only dysregulation of m6A-machinery genes is detectable in multiple brain regions and in the periphery, but altered expression of m6A-methylome is affected by the number of m6A modifications peaks on the genes.

## Conclusions & Future Directions

The field of epitranscriptomics is rapidly evolving and largely driven by advancements in massively parallel sequencing, bioinformatics, and public databases. Despite the widespread nature of m6A modification sites, the majority are unmethylated at baseline ^121^. The functional relevance of the constitutive m6A sites versus regulated m6A sites is unknown but such sub-stoichiometric levels of m6A would indicate a large margin in which to regulate neuronal gene expression. Furthermore, while the lifecycle of m6A modifications is thought to be dynamic, the degree of dynamicity and whether it varies from transcript to transcript is unknown. The apparent clustering of modifications on the same transcript also suggests that individual modifications are unlikely to have a significant functional effect on their own; rather they may interact with each other in as yet unknown ways to influence transcript stability and gene expression as can be seen in our cumulative distribution plot analyses. Adding to this complexity is our observation that there does not appear to be a clear correlation between the number of m6A sites and differential gene expression. Therefore, to functionally interrogate specific modification sites is challenging. Accordingly, using strategies to manipulate the epitranscriptomic machinery of ‘writers’, ‘erasers’ and ‘readers’ while low in resolution, is relatively easier and just as informative in the interim, until RNA-modification detection, mapping and manipulation technologies have been refined and matured. Additionally, future investigations should consider incorporating techniques such as ribosome profiling and proteomics (Buccitelli & Selbach, 2020).

In terms of the mechanisms which link stress exposure to changes in the m6A-methylome and/or functional expression of m6A machinery, we speculate that they are likely embedded in the wave of responses downstream of CRFR, GR or MR activation ^98,122,^ for reviews see ^123,124^. For example, this could include the transport and ‘on-demand’ processing of m6A-RNAs (e.g. FAAH) in distal compartments such as the synapse. Additionally, stress can induce expression of non-coding RNAs which are susceptible to m6A-methylation, thus interfering with or enhancing their ability to finetune expression of genes which may include those involved in the stress response or even which form the m6A-machinery. Taken together, this array of complex transcriptional and post-transcriptional effects may underpin, in part, changes in synaptic plasticity, neurotransmission and energy metabolism required to mount a transient, behavioural response to stress that is adaptive. Maladaptive responses may emerge if any of the above mechanisms are not efficiently terminated, leading to the onset of behavioural phenotypes of relevance to major depression and PTSD. In this regard, research into the role of the m6A-methylome/machinery as potential mediators of resilience to stress is an exciting prospect.

As we have shown, very few human studies have been carried out on the epitranscriptome, but our analysis of publicly available data shows sex-differences in major depression and PTSD. Future research should also consider the potential influence of individual risk factors (e.g. early-life adversity, drug abuse) on the m6A-methylome/machinery and whether these interactions influence disorder onset. It is worth highlighting that given the clinical presentation of these disorders is heterogenous and varies with time, we have yet to explore the short- and long-term impact of stress on the m6A-methylome/machinery and how they might correlate with clinically relevant phenotypes.

And finally, evidence has shown that post-transcriptional mechanisms are implicated in the mechanism of action for therapeutics for the treatment of major depression and PTSD (O’Connor *et al.*, 2012; Murphy & Singewald, 2018). It would be interesting to examine if m6A-methylome/machinery is similarly amenable to therapeutic targeting.

In conclusion, our review has highlighted significant findings from animal and human studies in support of a hypothesis for the involvement of the m6A-methylome/machinery in the pathophysiology of stress-induced disorders such as major depression and PTSD. Future research incorporating the experimental strategies described above will significantly advance our understanding on the impact of stress on epitranscriptomic mechanisms in the brain in both healthy and psychiatric states. These mechanisms may in turn form the basis for developing next-generation therapeutics to treat these disorders.

## Acknowledgements

DOW is supported by KAKENHI 17H03546, 19H04907, 19H05212, AMED 18dm0307023h 0001, NSFC 31971335, Xingliao Talents Program, XLYC1802007, Department of Education of Liaoning Province 1911520092, Hirose research grant, Takeda research grant. AG is supported by University of Sydney Research Fellowship.

