## Supplementary Methods, Figures and Tables for "Epitranscriptomic Dysregulation in Stress-induced Psychopathologies"

### Supplementary Information

**Supplementary Table 1**

This data in this table was generated using miRNET 2.0^1^. The m6A-machinery genes were entered as ‘Genes’ into a query list. The organism specified was ‘H.Sapiens (human) and ‘Brain’ was selected for tissue type. We selected ‘miRNA’ for the ‘Target by’ option.

| ID | Accession | Target | TargetID | Experiment | Literature | Tissue |
| --- | --- | --- | --- | --- | --- | --- |
| hsa-mir-193a-3p | MIMAT0000459 | ALKBH5 | 54890 | HITS-CLIP | 22473208 | Brain |
| hsa-mir-125a-3p | MIMAT0004602 | ALKBH5 | 54890 | PAR-CLIP | 23592263\|26701625 | Brain |
| hsa-mir-186-3p | MIMAT0004612 | ALKBH5 | 54890 | HITS-CLIP | tarbase | Brain |
| hsa-mir-186-5p | MIMAT0000456 | ALKBH5 | 54890 | PAR-CLIP | tarbase | Brain |
| hsa-mir-218-5p | MIMAT0000275 | ALKBH5 | 54890 | PAR-CLIP | tarbase | Brain |
| hsa-mir-377-3p | MIMAT0000730 | ALKBH5 | 54890 | HITS-CLIP | tarbase | Brain |
| hsa-mir-708-5p | MIMAT0004926 | ALKBH5 | 54890 | HITS-CLIP | tarbase | Brain |
| hsa-mir-1-3p | MIMAT0000416 | ALKBH5 | 54890 | RPF-Seq | tarbase | Brain |
| hsa-mir-155-5p | MIMAT0000646 | ALKBH5 | 54890 | Microarrays | tarbase | Brain |
| hsa-mir-16-5p | MIMAT0000069 | ALKBH5 | 54890 | Microarrays | tarbase | Brain |
| hsa-let-7b-5p | MIMAT0000063 | FTO | 79068 | CLASH | 23622248 | Brain |
| hsa-mir-143-5p | MIMAT0004599 | FTO | 79068 | HITS-CLIP | tarbase | Brain |
| hsa-mir-33a-5p | MIMAT0000091 | FTO | 79068 | PAR-CLIP | tarbase | Brain |
| hsa-mir-1-3p | MIMAT0000416 | FTO | 79068 | RPF-Seq, RNA-Seq | tarbase | Brain |
| hsa-mir-16-5p | MIMAT0000069 | FTO | 79068 | Microarrays | tarbase | Brain |
| hsa-mir-149-3p | MIMAT0004609 | METTL14 | 57721 | PAR-CLIP | 23446348 | Brain |
| hsa-mir-101-3p | MIMAT0000099 | METTL14 | 57721 | PAR-CLIP, HITS-CLIP | tarbase | Brain |
| hsa-mir-17-5p | MIMAT0000070 | METTL14 | 57721 | PAR-CLIP | tarbase | Brain |
| hsa-mir-186-5p | MIMAT0000456 | METTL14 | 57721 | PAR-CLIP | tarbase | Brain |
| hsa-mir-33a-5p | MIMAT0000091 | METTL14 | 57721 | PAR-CLIP | tarbase | Brain |
| hsa-mir-361-5p | MIMAT0000703 | METTL14 | 57721 | PAR-CLIP | tarbase | Brain |
| hsa-mir-93-5p | MIMAT0000093 | METTL14 | 57721 | PAR-CLIP | tarbase | Brain |
| hsa-mir-1-3p | MIMAT0000416 | METTL14 | 57721 | Microarrays | tarbase | Brain |
| hsa-mir-33a-3p | MIMAT0004506 | METTL3 | 56339 | Luciferase reporter assay//qRT-PCR//Western blot | 27856248 | Brain |
| hsa-mir-101-3p | MIMAT0000099 | METTL3 | 56339 | PAR-CLIP | tarbase | Brain |
| hsa-mir-186-5p | MIMAT0000456 | METTL3 | 56339 | PAR-CLIP | tarbase | Brain |
| hsa-let-7b-5p | MIMAT0000063 | YTHDC1 | 91746 | CLASH | 23622248 | Brain |
| hsa-mir-15a-5p | MIMAT0000068 | YTHDC1 | 91746 | PAR-CLIP | 23446348\|21572407\|20371350 | Brain |
| hsa-mir-16-5p | MIMAT0000069 | YTHDC1 | 91746 | PAR-CLIP//Sequencing | 20371350\|23446348\|21572407 | Brain |
| hsa-mir-17-5p | MIMAT0000070 | YTHDC1 | 91746 | HITS-CLIP | 19536157 | Brain |
| hsa-mir-24-3p | MIMAT0000080 | YTHDC1 | 91746 | HITS-CLIP | 23824327 | Brain |
| hsa-mir-93-5p | MIMAT0000093 | YTHDC1 | 91746 | HITS-CLIP//Sequencing | 20371350\|19536157 | Brain |
| hsa-mir-107 | MIMAT0000104 | YTHDC1 | 91746 | PAR-CLIP | 23446348\|21572407\|20371350 | Brain |
| hsa-mir-1-3p | MIMAT0000416 | YTHDC1 | 91746 | Proteomics | 18668040 | Brain |
| hsa-mir-30e-3p | MIMAT0000693 | YTHDC1 | 91746 | HITS-CLIP | 19536157 | Brain |
| hsa-mir-20a-3p | MIMAT0004493 | YTHDC1 | 91746 | CLASH | 23622248 | Brain |
| hsa-mir-214-5p | MIMAT0004564 | YTHDC1 | 91746 | HITS-CLIP | 19536157 | Brain |
| hsa-mir-143-5p | MIMAT0004599 | YTHDC1 | 91746 | HITS-CLIP | 23824327 | Brain |
| hsa-mir-95-5p | MIMAT0026473 | YTHDC1 | 91746 | PAR-CLIP | 21572407 | Brain |
| hsa-mir-101-3p | MIMAT0000099 | YTHDC1 | 91746 | PAR-CLIP | tarbase | Brain |
| hsa-mir-101-3p | MIMAT0000099 | YTHDC1 | 91746 | PAR-CLIP | tarbase | Brain |
| hsa-mir-129-5p | MIMAT0000242 | YTHDC1 | 91746 | HITS-CLIP | tarbase | Brain |
| hsa-mir-129-5p | MIMAT0000242 | YTHDC1 | 91746 | HITS-CLIP | tarbase | Brain |
| hsa-mir-143-3p | MIMAT0000435 | YTHDC1 | 91746 | PAR-CLIP | tarbase | Brain |
| hsa-mir-143-3p | MIMAT0000435 | YTHDC1 | 91746 | PAR-CLIP | tarbase | Brain |
| hsa-mir-204-5p | MIMAT0000265 | YTHDC1 | 91746 | HITS-CLIP | tarbase | Brain |
| hsa-mir-204-5p | MIMAT0000265 | YTHDC1 | 91746 | HITS-CLIP | tarbase | Brain |
| hsa-mir-212-5p | MIMAT0022695 | YTHDC1 | 91746 | HITS-CLIP | tarbase | Brain |
| hsa-mir-212-5p | MIMAT0022695 | YTHDC1 | 91746 | HITS-CLIP | tarbase | Brain |
| hsa-mir-191-5p | MIMAT0000440 | YTHDC1 | 91746 | Microarrays | tarbase | Brain |
| hsa-mir-191-5p | MIMAT0000440 | YTHDC1 | 91746 | Microarrays | tarbase | Brain |
| hsa-let-7a-5p | MIMAT0000062 | YTHDC2 | 64848 | PAR-CLIP | tarbase | Brain |
| hsa-let-7b-5p | MIMAT0000063 | YTHDC2 | 64848 | PAR-CLIP | tarbase | Brain |
| hsa-mir-128-3p | MIMAT0000424 | YTHDC2 | 64848 | HITS-CLIP | tarbase | Brain |
| hsa-mir-21-5p | MIMAT0000076 | YTHDC2 | 64848 | Microarrays | tarbase | Brain |
| hsa-mir-23a-3p | MIMAT0000078 | YTHDC2 | 64848 | HITS-CLIP | tarbase | Brain |
| hsa-mir-23b-3p | MIMAT0000418 | YTHDC2 | 64848 | HITS-CLIP | tarbase | Brain |
| hsa-mir-137 | MIMAT0000429 | YTHDC2 | 64848 | HITS-CLIP | tarbase | Brain |
| hsa-mir-1-3p | MIMAT0000416 | YTHDC2 | 64848 | Microarrays | tarbase | Brain |
| hsa-mir-139-5p | MIMAT0000250 | YTHDF1 | 54915 | PAR-CLIP | 20371350 | Brain |
| hsa-mir-218-5p | MIMAT0000275 | YTHDF1 | 54915 | HITS-CLIP | 23212916 | Brain |
| hsa-mir-149-5p | MIMAT0000450 | YTHDF1 | 54915 | CLASH | 23622248 | Brain |
| hsa-mir-186-5p | MIMAT0000456 | YTHDF1 | 54915 | PAR-CLIP | 21572407 | Brain |
| hsa-mir-612 | MIMAT0003280 | YTHDF1 | 54915 | PAR-CLIP | 20371350 | Brain |
| hsa-mir-767-3p | MIMAT0003883 | YTHDF1 | 54915 | HITS-CLIP | 23313552 | Brain |
| hsa-let-7a-5p | MIMAT0000062 | YTHDF1 | 54915 | PAR-CLIP | tarbase | Brain |
| hsa-let-7b-5p | MIMAT0000063 | YTHDF1 | 54915 | PAR-CLIP | tarbase | Brain |
| hsa-mir-107 | MIMAT0000104 | YTHDF1 | 54915 | HITS-CLIP, PAR-CLIP | tarbase | Brain |
| hsa-mir-124-3p | MIMAT0000422 | YTHDF1 | 54915 | HITS-CLIP, Microarrays | tarbase | Brain |
| hsa-mir-15a-5p | MIMAT0000068 | YTHDF1 | 54915 | HITS-CLIP | tarbase | Brain |
| hsa-mir-16-5p | MIMAT0000069 | YTHDF1 | 54915 | HITS-CLIP, PAR-CLIP | tarbase | Brain |
| hsa-mir-17-5p | MIMAT0000070 | YTHDF1 | 54915 | PAR-CLIP, HITS-CLIP | tarbase | Brain |
| hsa-mir-18a-5p | MIMAT0000072 | YTHDF1 | 54915 | PAR-CLIP, HITS-CLIP | tarbase | Brain |
| hsa-mir-23a-3p | MIMAT0000078 | YTHDF1 | 54915 | HITS-CLIP, PAR-CLIP | tarbase | Brain |
| hsa-mir-23b-3p | MIMAT0000418 | YTHDF1 | 54915 | HITS-CLIP, Microarrays | tarbase | Brain |
| hsa-mir-24-3p | MIMAT0000080 | YTHDF1 | 54915 | HITS-CLIP | tarbase | Brain |
| hsa-mir-29a-3p | MIMAT0000086 | YTHDF1 | 54915 | HITS-CLIP | tarbase | Brain |
| hsa-mir-30e-3p | MIMAT0000693 | YTHDF1 | 54915 | HITS-CLIP | tarbase | Brain |
| hsa-mir-342-3p | MIMAT0000753 | YTHDF1 | 54915 | HITS-CLIP | tarbase | Brain |
| hsa-mir-361-5p | MIMAT0000703 | YTHDF1 | 54915 | HITS-CLIP | tarbase | Brain |
| hsa-mir-383-5p | MIMAT0000738 | YTHDF1 | 54915 | HITS-CLIP | tarbase | Brain |
| hsa-mir-603 | MIMAT0003271 | YTHDF1 | 54915 | HITS-CLIP | tarbase | Brain |
| hsa-mir-93-5p | MIMAT0000093 | YTHDF1 | 54915 | PAR-CLIP, HITS-CLIP | tarbase | Brain |
| hsa-mir-95-3p | MIMAT0000094 | YTHDF1 | 54915 | HITS-CLIP, Microarrays | tarbase | Brain |
| hsa-mir-99b-3p | MIMAT0004678 | YTHDF1 | 54915 | HITS-CLIP | tarbase | Brain |
| hsa-mir-1-3p | MIMAT0000416 | YTHDF1 | 54915 | RPF-Seq | tarbase | Brain |
| hsa-mir-191-5p | MIMAT0000440 | YTHDF1 | 54915 | Microarrays | tarbase | Brain |
| hsa-mir-221-3p | MIMAT0000278 | YTHDF1 | 54915 | Chimeric fragments | tarbase | Brain |
| hsa-mir-324-5p | MIMAT0000761 | YTHDF1 | 54915 | Chimeric fragments | tarbase | Brain |
| hsa-mir-1-3p | MIMAT0000416 | YTHDF2 | 51441 | Proteomics | 18668040 | Brain |
| hsa-mir-107 | MIMAT0000104 | YTHDF2 | 51441 | HITS-CLIP | tarbase | Brain |
| hsa-mir-124-5p | MIMAT0004591 | YTHDF2 | 51441 | HITS-CLIP | tarbase | Brain |
| hsa-mir-125a-5p | MIMAT0000443 | YTHDF2 | 51441 | HITS-CLIP | tarbase | Brain |
| hsa-mir-125b-5p | MIMAT0000423 | YTHDF2 | 51441 | HITS-CLIP | tarbase | Brain |
| hsa-mir-138-5p | MIMAT0000430 | YTHDF2 | 51441 | Microarrays | tarbase | Brain |
| hsa-mir-153-3p | MIMAT0000439 | YTHDF2 | 51441 | HITS-CLIP | tarbase | Brain |
| hsa-mir-186-3p | MIMAT0004612 | YTHDF2 | 51441 | PAR-CLIP | tarbase | Brain |
| hsa-mir-186-5p | MIMAT0000456 | YTHDF2 | 51441 | PAR-CLIP | tarbase | Brain |
| hsa-mir-30d-5p | MIMAT0000245 | YTHDF2 | 51441 | HITS-CLIP | tarbase | Brain |
| hsa-mir-33a-5p | MIMAT0000091 | YTHDF2 | 51441 | HITS-CLIP, PAR-CLIP | tarbase | Brain |
| hsa-mir-124-3p | MIMAT0000422 | YTHDF2 | 51441 | Microarrays | tarbase | Brain |
| hsa-mir-191-5p | MIMAT0000440 | YTHDF2 | 51441 | Microarrays | tarbase | Brain |
| hsa-mir-23b-3p | MIMAT0000418 | YTHDF2 | 51441 | Microarrays | tarbase | Brain |
| hsa-mir-7-5p | MIMAT0000252 | YTHDF2 | 51441 | Microarrays | tarbase | Brain |
| hsa-mir-137 | MIMAT0000429 | YTHDF3 | 253943 | HITS-CLIP | 19536157 | Brain |
| hsa-mir-124-5p | MIMAT0004591 | YTHDF3 | 253943 | PAR-CLIP | 20371350 | Brain |
| hsa-let-7a-5p | MIMAT0000062 | YTHDF3 | 253943 | HITS-CLIP, Chimeric fragments | tarbase | Brain |
| hsa-let-7b-5p | MIMAT0000063 | YTHDF3 | 253943 | HITS-CLIP, Chimeric fragments | tarbase | Brain |
| hsa-mir-125a-5p | MIMAT0000443 | YTHDF3 | 253943 | HITS-CLIP, PAR-CLIP | tarbase | Brain |
| hsa-mir-125b-5p | MIMAT0000423 | YTHDF3 | 253943 | HITS-CLIP, PAR-CLIP | tarbase | Brain |
| hsa-mir-139-5p | MIMAT0000250 | YTHDF3 | 253943 | PAR-CLIP | tarbase | Brain |
| hsa-mir-141-3p | MIMAT0000432 | YTHDF3 | 253943 | HITS-CLIP | tarbase | Brain |
| hsa-mir-17-5p | MIMAT0000070 | YTHDF3 | 253943 | PAR-CLIP, HITS-CLIP | tarbase | Brain |
| hsa-mir-181b-5p | MIMAT0000257 | YTHDF3 | 253943 | PAR-CLIP | tarbase | Brain |
| hsa-mir-186-5p | MIMAT0000456 | YTHDF3 | 253943 | HITS-CLIP | tarbase | Brain |
| hsa-mir-18a-5p | MIMAT0000072 | YTHDF3 | 253943 | HITS-CLIP | tarbase | Brain |
| hsa-mir-199a-3p | MIMAT0000232 | YTHDF3 | 253943 | PAR-CLIP | tarbase | Brain |
| hsa-mir-200a-3p | MIMAT0000682 | YTHDF3 | 253943 | HITS-CLIP | tarbase | Brain |
| hsa-mir-212-3p | MIMAT0000269 | YTHDF3 | 253943 | HITS-CLIP | tarbase | Brain |
| hsa-mir-30d-5p | MIMAT0000245 | YTHDF3 | 253943 | PAR-CLIP | tarbase | Brain |
| hsa-mir-30e-3p | MIMAT0000693 | YTHDF3 | 253943 | HITS-CLIP, PAR-CLIP | tarbase | Brain |
| hsa-mir-33a-5p | MIMAT0000091 | YTHDF3 | 253943 | HITS-CLIP | tarbase | Brain |
| hsa-mir-93-3p | MIMAT0004509 | YTHDF3 | 253943 | PAR-CLIP | tarbase | Brain |
| hsa-mir-93-5p | MIMAT0000093 | YTHDF3 | 253943 | PAR-CLIP, HITS-CLIP | tarbase | Brain |
| hsa-mir-9-5p | MIMAT0000441 | YTHDF3 | 253943 | PAR-CLIP, HITS-CLIP | tarbase | Brain |
| hsa-let-7e-5p | MIMAT0000066 | YTHDF3 | 253943 | HITS-CLIP | tarbase | Brain |
| hsa-mir-1-3p | MIMAT0000416 | YTHDF3 | 253943 | RPF-Seq | tarbase | Brain |
| hsa-mir-191-5p | MIMAT0000440 | YTHDF3 | 253943 | RIP-Seq | tarbase | Brain |
| hsa-mir-494-3p | MIMAT0002816 | YTHDF3 | 253943 | RIP-Seq, RNA-Seq | tarbase | Brain |
| hsa-mir-124-3p | MIMAT0000422 | YTHDF3 | 253943 | Other | tarbase | Brain |
| hsa-mir-155-5p | MIMAT0000646 | YTHDF3 | 253943 | Chimeric fragments | tarbase | Brain |
| hsa-mir-222-3p | MIMAT0000279 | YTHDF3 | 253943 | Chimeric fragments | tarbase | Brain |

##### Datasets represented in Figures 3, 4, 5, 6, 7 and Fig S1

RNA-seq and microarray expression profiling datasets related to case and control studies for major depression and PTSD) were obtained from the NCBI GEO database (Supplementary Table 2).

**Supplementary Table 2**

| NCBI Geo Accession ID | Study title |
| --- | --- |
| GSE144136 | Single-nucleus RNA-seq in the post-mortem brain in major depressive disorder |
| GSE54571 | Expression data from human brain anterior cingulate cortex - including control samples and samples with major depression disorders (26 samples BA25_F) |
| GSE54572 | Expression data from human brain anterior cingulate cortex - including control samples and samples with major depression disorders (24 samples BA25_M) |
| GSE80655 | RNA-sequencing of human post-mortem brain tissues |
| GSE54565 | Expression data from human brain anterior cingulate cortex - including control samples and samples with major depression disorders (32 samples MD1_ACC) |
| GSE54563 | Expression data from human brain anterior cingulate cortex - including control samples and samples with major depression disorders (50 samples MD3_ACC) |
| GSE54562 | Expression data from human brain anterior cingulate cortex - including control samples and samples with major depression disorders (20 samples MD2_ACC) |
| GSE44593 | Molecular Evidence for a Dimensional Basis of Depression |
| GSE54566 | Expression data from human brain amygdala - including control samples and samples with major depression disorders (28 samples MD1_AMY) |
| GSE54564 | Expression data from human brain amygdala - including control samples and samples with major depression disorders (42 samples MD3_AMY) |
| GSE35974 | Expression data from the human cerebellum brain |
| GSE54568 | Expression data from human brain dorsolateral prefrontal cortex - including control samples and samples with major depression disorders (30 samples BA9_F) |
| GSE54567 | Expression data from human brain dorsolateral prefrontal cortex - including control samples and samples with major depression disorders (28 samples BA9_M) |
| GSE101521 | Whole-transcriptome brain expression and exon-usage profiling in major depression and suicide |
| GSE54570 | Expression data from human brain dorsolateral prefrontal cortex - including control samples and samples with major depression disorders (26 samples NY_BA9) |
| GSE87610 | Gene expression of L3 and L5 pyramidal neurons in the DLPFC comparing schizophrenia from bipolar major depressive disorders and unaffected subjects. |
| GSE92538 | Inference of cell-type composition from human brain transcriptomic datasets illuminates the effects of age, manner of death, dissection, and psychiatric diagnosis |
| GSE17440 | Gene Expression in Frontal Cortex in Major Depression and HIV |
| GSE24095 | Human postmortem hippocampus: Major depressive disorder (MDD) vs. Control |
| GSE53987 | Microarray profiling of PFC, HPC and STR from subjects with schizophrenia, bipolar, MDD or control |
| GSE54575 | Expression data from human brain orbital ventral prefrontal cortex - including control samples and samples with major depression disorders (24 samples NY_BA47) |
| GSE35977 | Expression data from the human parietal cortex brain |
| GSE12654 | Gene expression from human prefrontal cortex (BA10) |
| GSE81761 | Gene Expression Pathways Implicated in Posttraumatic Stress Disorder and Symptomatic Improvement |
| GSE77164 | Peripheral blood transcriptome profiles in Nepali child soldiers and civilians. |
| GSE67663 | Gene expression profiles in comorbid PTSD and depression |
| GSE860 | PTSD Normalized data |
| GSE109409 | Using Next-Generation Sequencing Transcriptomics to Determine Markers of Post-traumatic Symptoms - preliminary findings from a post-deployment cohort |
| GSE97356 | Gene expression associated with PTSD in World Trade Center responders: An RNA sequencing study |
| GSE83601 | Dysregulated immune system networks in war veterans with PTSD |

##### Differential expression analysis of snRNA-seq

Data was obtained from GSE144136. Cell ranger output was imported as a cell matrix using the *Seurat* package and converted to a *Seurat* object in R. The counts are normalized using the *NormalizeData* function from the *Seurat* package. Cell types, which were predefined by the authors, were assigned to the features. Mean log-normalised expression of the genes associated with the m6A machinery were also plotted as heatmap. Differential expression of genes within each cell type between cases and controls was determined using *FindMarkers* function with a p-value cut-off < 0.05.

##### Differential expression analysis of microarray and RNAseq datasets

For microarray datasets, webtools GEO2R was used for performing differential expression of genes. Briefly, datasets were imported to GEO2R from GEO database. User-defined groups (i.e. case and control) were assigned to individual samples. The samples were compared for differential expression of the genes. GEO2R uses eBayes function from the limma package to calculate significance of differential expression of genes between groups. Significant differential genes were selected based on p value cut-off < 0.05.

For RNA-seq datasets, total counts for genes were downloaded and normalized using *calcnormfactor* function from edgeR package followed by significance testing for differential expression using eBayes function from the limma package. For Jaffe et al.^2^, logFC and p values were precalculated by the authors using the limma package for different factors and conditions. We used these values directly for the investigation and plotting. Significant differential genes are selected based on p value cut-off < 0.05.

**Cumulative distribution of differentially expressed genes for methylated vs non-methylated genes**

The number of methylation sites on each gene was calculated from MeRIP-seq datasets (Jun’e Liu et.al. Mol Cell 2020) from post-mortem human cerebrum and cerebellum of two individuals. Raw reads were downloaded and quality trimmed to include high quality reads by *TrimGalore*. Trimmed reads were aligned to the human reference genome (hg38) using splice aware aligner *STAR*. We then used *exomePeak* with default parameters to identify m6a peaks in cerebrum and cerebellum samples. For each study, the differentially expressed genes were divided into 4 main groups based on the number of m6a sites mapped to their RNA. m6a_0: no m6a peaks; m6a_1: one m6a peak; m6a_2_5: two to five m6a peaks; m6a_6: six or more m6a peaks. DLPFC: dorsolateral prefrontal cortex; ACC: anterior cingulate cortex. Cumulative distribution function (CDF) plot is generated for four groups for each study using R.

Custom R scripts are available at https://github.com/kandarpRJ/epi_psych_metaanalysis.

**Supplementary Figure 1**

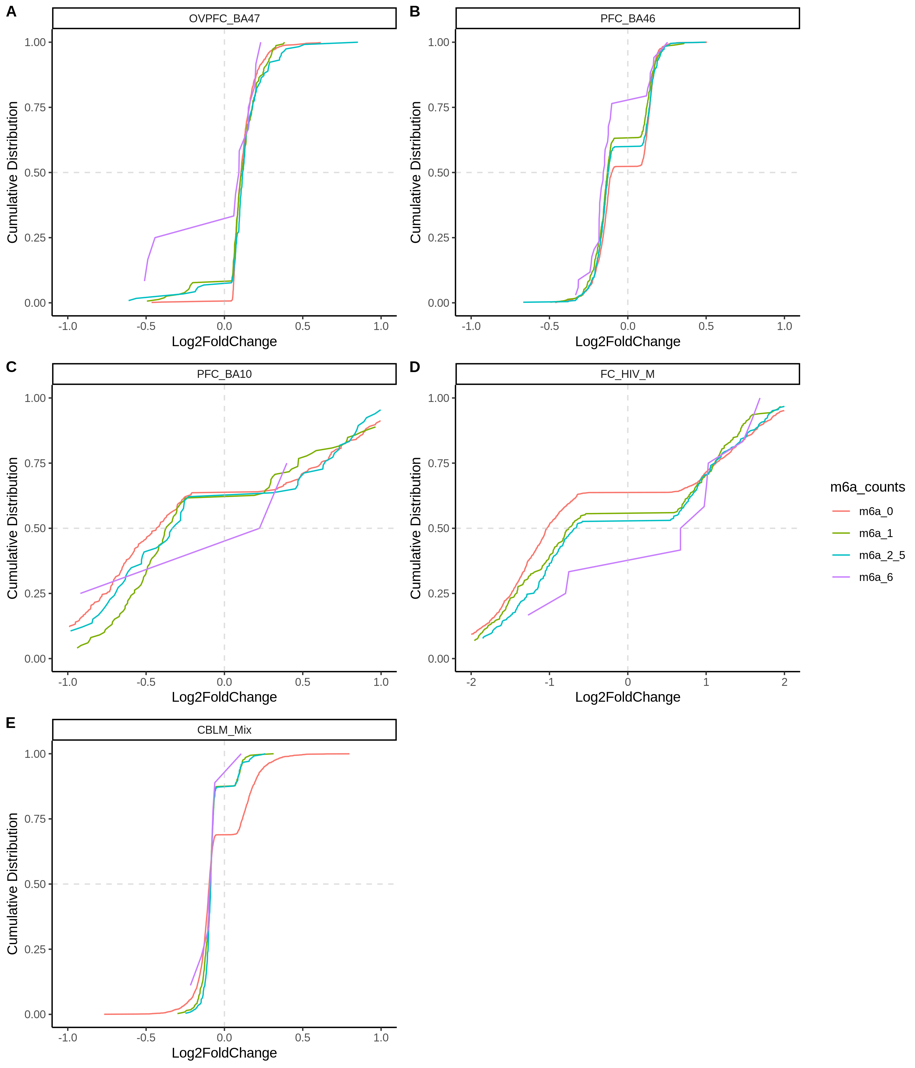

Figure S1 - Cumulative distributions of log2FC of gene expression in MDD patients vs control. The differentially expressed genes are divided into four groups based on the number of m6a sites mapped to their RNA. m6a_0: no m6a peaks; m6a_1: one m6a peak; m6a_2_5: two to five m6a peaks; m6a_6: six or more m6a peaks. OVPFC: orbital ventral prefrontal cortex; PFC: prefrontal cortex; FC: frontal cortex; ; CBLM: cerebellum.

Analysis of samples derived from orbital ventral prefrontal cortex (OVPFC_BA47) revealed an overall greater proportion of upregulated genes; for genes with more than 6 m6A sites, this was less (Supplementary Figure 1A). In PFC_BA46 there was a greater proportion of downreguled of genes having six or more m6A sites as compared to other groups (Figure B). In PFC_BA10, genes with five or less m6A sites showed more downregulation in expression as compared to genes with more than 6 m6A sites (Supplementary Figure 1C) ^3^.

In a study which examined samples from the frontal cortex of male HIV patients (FC_HIV_M) who had suffered from depression, we observed a higher degree of overall dysregulation in gene (Supplementary Figure 1D). For genes with no m6A sites, the proportion of downregulated genes was higher whereas for genes with six or more m6A sites, the proportion of upregulated genes was higher ^4^.

Finally, we also performed similar analysis in datasets generated from the cerebellum of patients and control subjects (CBLM_Mix)^5^. We observed a greater proportion of downregulated overall with genes having m6A sites more down regulated than genes with no m6A sites as observed by leftward shift of cumulative distribution (Supplementary Figure 1E).
